# Comprehensive analysis of the effect of mRNA sequences on translation efficiency and accuracy

**DOI:** 10.1101/2022.05.19.492606

**Authors:** S. Umemoto, T. Kondo, T. Fujino, G. Hayashi, H. Murakami

## Abstract

Messenger ribonucleic acid (mRNA) sequences influence the translation efficiency and accuracy. To increase our knowledge of how mRNA sequences affect ribosome translation and apply the obtained information to improve the mRNA display method, we conducted a comprehensive analysis of the effect of mRNA sequences on the translation. Translation efficiency depended strongly on the three codons following the start codon. Furthermore, the codons at the ribosomal E- and P-sites strongly influence the misreading of the A-site blank codon by near-cognate transfer RNA. The purine base after the blank codon also induced a higher misread rate than that with a pyrimidine base. Based on these findings, we demonstrated construction of highly diverse monobody and macrocyclic peptide libraries that would be useful in developing functional peptides and proteins in the future.

## Introduction

Ribosomal translation efficiency and accuracy are evolutionally optimized to maintain protein production and function. Translation efficiency is mainly affected by translation initiation and early elongation of a protein involving the messenger ribonucleic acid (mRNA) secondary structure^1-4^, AT richness^2,5^, transfer RNA (tRNA) abundance^6,7^, and positively charged amino acids^8^. The stable mRNA secondary structure near the start codon affects decoding protein expression by decreasing translation initiation efficiency^4^. In bacteria, a high AT content at the 5′ end of the open reading frame (ORF) is correlated with higher protein expression^2,5^. The codons cognate to rare tRNAs^7^ and those encoding positively charged amino acids^8^ are enriched at the *N*-terminus of genes in bacteria.

Large-scale analyses of correlations between mRNA sequences of the *N*-terminal region and green fluorescent protein (GFP) production using fluorescence-activated cell sorting (FACS) followed by next-generation sequencing (NGS) indicate that stability of the mRNA structure near the start codon affects translation efficiency^9-13^. AT richness and rare codons accumulates at the mRNA of the *N*-terminal region to increase translation efficiency by forming a weak mRNA secondary structure^10^. These reports showed that mRNA sequences of the protein *N*-terminal region play an important role in determining translation efficiency. However, the previous *in vivo* studies on translation potentially involved the mRNA transcription efficiency and stability and the GFP folding efficiency.

Translation accuracy depends on the presence or absence of tRNA at the E-site of the ribosome^14-16^. Ribosomes with cognate tRNAs at the E-site have low-affinity and high-fidelity A-sites that effectively distinguish between cognate and noncognate tRNAs. In contrast, ribosomes with empty E-sites have high-affinity and low-fidelity A-sites. This finding is consistent with the observation that the activation energy for A-site binding is greater when the E-site is occupied by a cognate tRNA^17^. Interestingly, Mottagui-Tabar et al. showed that the E-site codon differently affects misreading of the UGA stop codon at the A-site by tRNA^Trp^ (near-cognate) *in vivo*^18^. This result indicates that the types of tRNA species at the ribosomal E-site might also affect translation accuracy. However, to the best of our knowledge, no study has presented a large-scale analysis of the effect of E- or P-site codons on translation accuracy.

This study directly and comprehensively investigated how the mRNA sequence affects translation efficiency and accuracy by using an mRNA display method involving an *Escherichia coli*-based reconstituted *in vitro* translation system. First, we comprehensively analyzed the effect of the three codons after the start codon on translation efficiency. An enrichment factor, introduced to evaluate translation efficiency, varied over a 40-fold range, depending on the three-codon sequences. Second, we comprehensively analyzed the effect of E- and P-site codons and the single base after the A-site codon on translation accuracy. The enrichment factor was used to evaluate the frequency of misreading the UAG blank codon at the A-site. The E- and P-site codons as well as the single base after the A-site codon independently affected the enrichment factor. Notably, the effects of mRNA sequences on translation accuracy were commonly observed in different coding proteins, suggesting the generalizability of the results.

## Results

### Method for comprehensive analysis of the effect of mRNA sequences on translation efficiency

The previous large-scale analyses utilized a bacterial expression system that may involve mRNA transcription efficiency and stability *in vivo*^9-13^. To directly investigate translation efficiency, we employed a reconstituted *in vitro* translation system with 37 purified proteins and ribosomes^19-23^. We used an mRNA display method^23-26^ to conduct a comprehensive analysis of large number of mRNA sequences.

We prepared an mRNA library (SD8-fibronectin) carrying three NNW codons (N represents a mixture of A, U, G, and C; W represents a mixture of A and U; 32,768 sequences in theory) after a short 5′-UTR (SD8: 5′-GGAUUAAGGAGGUGAUAUUU-3′) and the start codon followed by the coding sequences of the 10th type III domain of human fibronectin (the backbone sequence of monobody^27^) and human influenza hemagglutinin-tag (HA-Tag) (Fig. 1a and Data file S1). The third base of codon was limited to W because these bases reportedly affected the translation efficiency positively^2,5^.

**Fig. 1.**
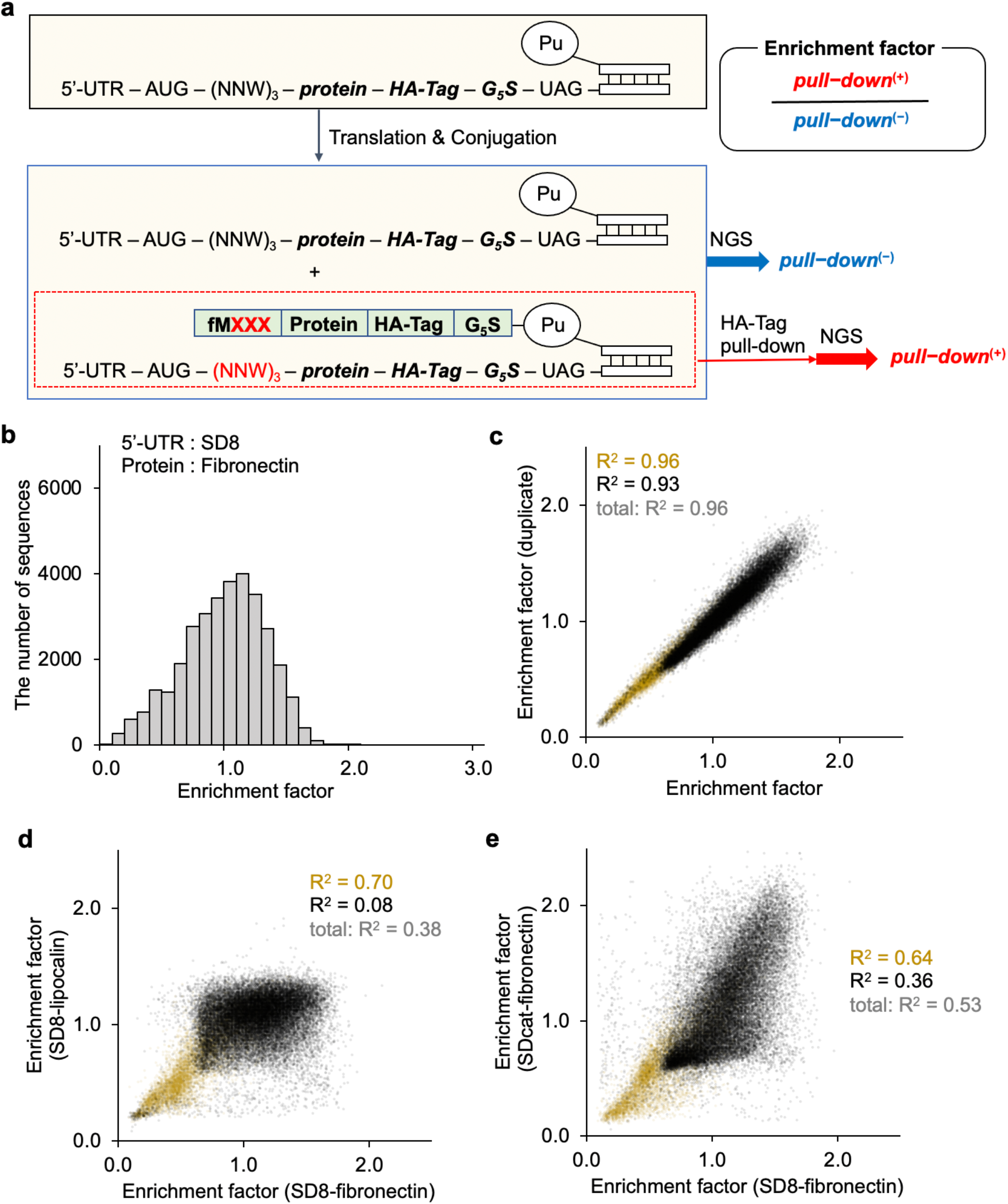
Comprehensive data sets of enrichment factors for mRNA libraries with AUG-NNW_3_ codons carrying the different 5′-UTRs and coding proteins to analyze the effect of the mRNA sequences on translation efficiency. **(a)** Method for analysis of the impact of mRNA sequences on translation efficiency. Three codons after the AUG start codon were randomized (AUG-NNW_3_; N = A, U, G, C; W = A, U), and different 5′-UTRs (SD8 and SDcat, see the main text for the sequences) and protein-coding sequences (the fibronectin and the lipocalin) were used in each mRNA library. The PuL-attached mRNA library was added to the RF-1 depleted translation system. Only the mRNAs with the coded protein were collected by pull-down via an anti-HA-Tag antibody. The enrichment factor was calculated from the ratio of the appearance rates before [*pull-down*^(−)^] and after the HA-Tag pull-down [*pull-down*^(+)^]. **(b)** Histogram of enrichment factors of all the sequences in the mRNA library encoding the SD8-fibronectin. Scatter plot of enrichment factors from **(c)** the duplicate experiments using the mRNA library encoding the SD8-fibronectin, **(d)** the SD8-fibronectin and SD8-lipocalin mRNA libraries, **(e)** the SD8-fibronectin and SDcat-fibronectin mRNA libraries. The yellow dots indicate the mRNAs carrying more than one stop codon or three consecutive Pro codons at the randomized codons. Abbreviations: Pu, puromycin; PuL, puromycin linker; 5′-UTR, 5′-untranslated region; SD, Shine–Dalgarno sequence; NGS, next-generation sequencing.

The mRNA library was annealed to a puromycin linker (PuL) and added to the cell-free translation system. In the translation system, ribosomes translate mRNAs to produce fibronectin and pause at the UAG codon after the G_5_S linker because the release factor 1 (RF-1) was excluded from the system. This effectively renders the UAG codon a blank. During the pause, the puromycin at the end of the mRNA attacks the ester bond of the peptidyl-tRNA, and the fibronectin-PuL/mRNA (“-” represents a covalent bond and “/” represents a noncovalent bond) complex is formed [*pull-down*^(-)^]. The resulting reaction mixture contained both translated and untranslated mRNAs, and only the translated mRNAs became fibronectin-PuL/mRNA complexes. To isolate translated mRNAs, we pulled down the fibronectin-PuL/mRNA complexes using the HA-Tag antibody [*pull-down*^(+)^]. The mRNA sequences in the *pull-down*^(-)^ and *pull-down*^(+)^ were analyzed by NGS. The appearance rate of each RNA sequence in *pull-down*^(+)^ was divided by the corresponding rate in *pull-down*^(-)^ to identify the enrichment factors. By principle, the enrichment factor reflected the translation efficiency of each mRNA carrying a unique sequence.

To see whether the protein-coding sequence and 5′-UTR influence the effect of the three codons on translation efficiency, we repeated the experiment via an mRNA library (SD8-lipocalin) in which the protein-coding sequence was replaced with human lipocalin-2 (the backbone of anticalin^28,29^) or SDcat-fibronectin whose 5′-UTR sequence was modified from SD8 to SDcat (5′-GGUUAAAGAGGAGAAAUUACAU-3′) (Data file S1).

### The effect of mRNA sequences near start codon on translation efficiency

We obtained more than 30 million NGS reads before and after pull-down in all three mRNA libraries (SD8-fibronectin, SD8-lipocalin, and SDcat-fibronectin) (Data file S2). The enrichment factor was distributed between 0.06 and 2.62, indicating that the three codons after the start codon significantly impacted translation efficiency (Figs. 1b and S1b). Replicating the experiment with an SD8-fibronectin mRNA library demonstrated data reliability (Fig. 1c, R^2^ = 0.96). As we expected, median enrichment factors of mRNAs carrying three-codon sequences with more than one stop codon were low in each library (0.52, SD8-fibronectin, Fig. 1c; 0.56, SD8-lipocalin Fig. 1d; 0.54, SDcat-fibronectin, Fig. 1e). Low median enrichment factors of mRNAs carrying consecutive three Pro (0.17, SD8-fibronectin; 0.26, SD8-lipocalin; 0.35, SDcat-fibronectin) were consistent with previous reports that the ribosome pauses at a Pro_3_ sequence in the absence of elongation factor P (EF-P)^30,31^ because of the unfavorable steric arrangement of peptidyl-Pro-tRNA^Pro^ in the P-site^32^. The effect of each three-codon sequence containing more than one stop codon or three consecutive Pro on the enrichment factors was independent of the protein-coding sequence and 5′-UTR sequence (yellow dots in Figs. 1d and 1e, R^2^ = 0.70 and 0.64) because the enrichment factors were reduced by same reasons.

The effect of the other sequences was strongly dependent on the protein-coding sequence (black dots in Fig. 1d, R^2^ = 0.08; the data from three-codon sequences with more than one stop codon or Pro_3_ were excluded to avoid bias). This contradicts prior reports showing that three kinds of *N*-terminal sequences influence protein expression levels regardless of the coding proteins^12^. The effect of each three-codon sequence on the enrichment factors was also dependent on the 5′-UTR sequence (black dots in Fig. 1e, R^2^ = 0.36), which again contradicted prior reports showing that three kinds of *N*-terminal sequences influence GFP expression levels regardless of 5′-UTR sequence^12^. These differences might be associated with differences between the *in vivo* and *in vitro* expression systems. Our expression system was constructed using only 37 kinds of purified proteins and ribosomes, thus, did not include various factors such as ribonuclease or proteases, which influence the stability of mRNAs and the produced proteins.

The sequences with a high enrichment factor (top 1%) in each mRNA library showed negligible base preference except for the third base of the codons (Fig. S2a). The average ratio of U and A at the third base (approximately NNU/NNA = 22/8) was greater than that in the whole mRNAs (NNU/NNA = 16/16). As the 2nd codon, CUU (Leu) and ACU (Thr) frequently appeared among all mRNA libraries (Fig. 2a). Pro and Leu codons were also commonly observed in all three positions (Fig. S2b).

**Fig. 2.**
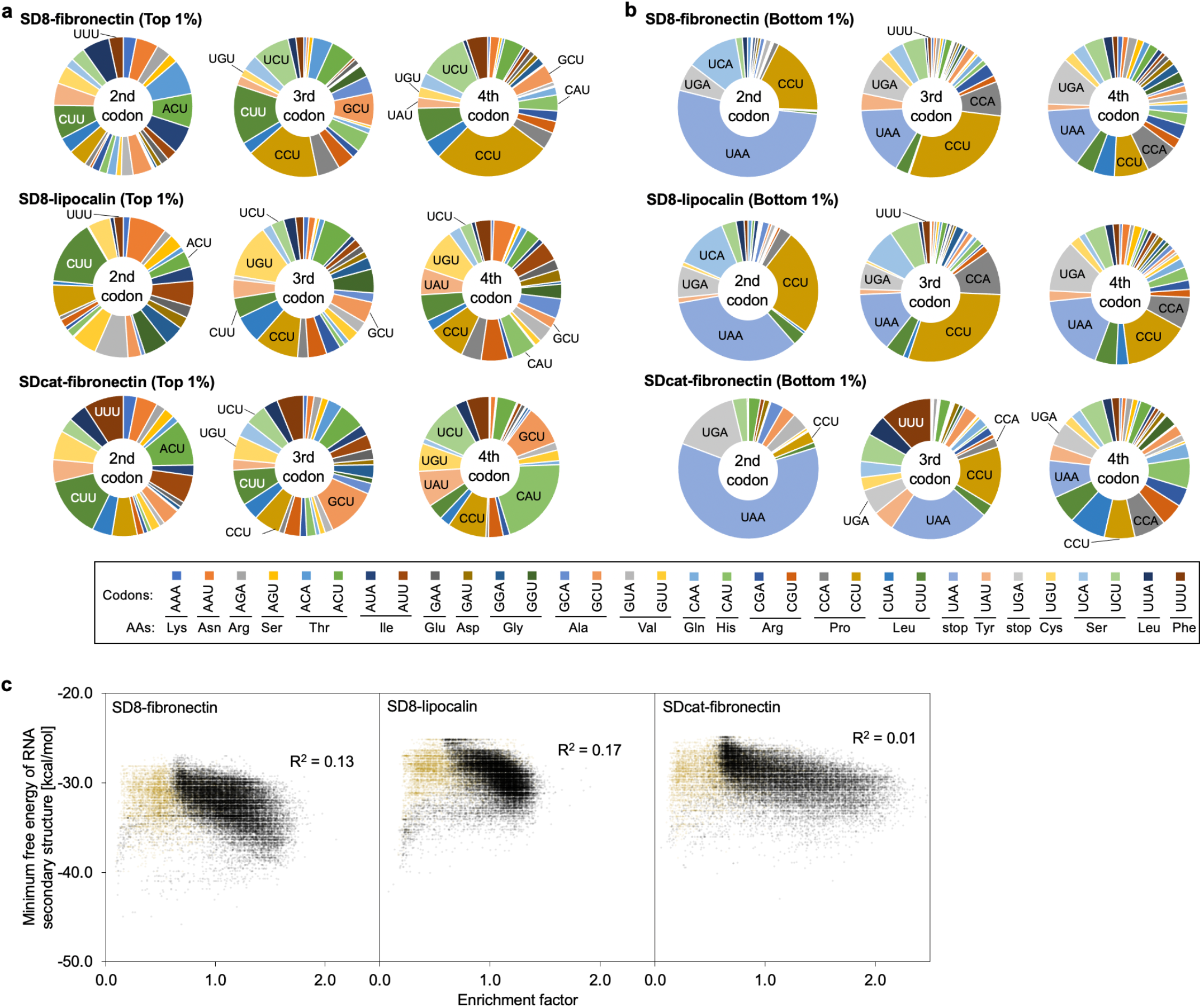
The effect of mRNA sequences on translation efficiency. (a) Circle graph of appearance frequencies of codons at each second, third, and fourth position in the mRNA having top 1% enrichment factors (327 sequences) and **(b)** those of bottom 1% enrichment factors (327 sequences). **(c)** Scatter plots between the enrichment factors and the minimum free energies of RNA secondary structure (5′-UTR followed by 112 bases of the protein-coding region) of all the sequences. The free energies were calculated using the RNAfold program from the Vienna RNA package^34^. The yellow dots indicate the mRNAs carrying more than one stop codon or three consecutive Pro codons at the randomized codons. Abbreviation: AA, amino acid. Other abbreviations were mentioned as previously.

The UAA and UGA stop codons frequently appeared in the sequences with a low enrichment factor (bottom 1%) for all three mRNA libraries (Fig. 2b). The frequencies in the second codon were higher than the ones in the third and fourth codons, suggesting that misreading was prevented in the second codon. This result was inconsistent with the fact that a ribosome with an empty E-site had a lower fidelity in an A-site than that with an occupied E-site^14,15^. The empty E-site might efficiently induce the peptidyl-tRNA drop-off during translocation or enhance RF-2 binding to the A-site, thus aborting peptide elongation. Three consecutive Lys also gave low enrichment factors in all mRNA libraries (0.57–0.74), whereas consecutive Arg showed moderate enrichment factors (0.54–1.63) (Figs. S2c–e). The poly-A sequence coding Lys induces sliding of the ribosome on the mRNA, causing frameshifts and premature termination^33^.

It is suggested that mRNA folding energy near the start codon determines protein expression level^1-4,9-13^. Therefore, we compared the enrichment factor and folding energy of each mRNA (5′-UTR–AUG-109 bases) calculated by the ViennaRNA^34^. Contrary to our expectations, we saw less correlation between the enrichment factor and mRNA folding energy (Fig. 2c; R^2^ = 0.13 for SD8-fibronectin, R^2^ = 0.17 for SD8-lipocalin, R^2^ = 0.01 for SDcat-fibronectin; the data from three-codon sequences with more than one stop codon or Pro_3_ were excluded to avoid bias). We also observed a lower difference in folding energy between synonymous codon variants and mean fold change in enrichment factors (Fig. S3a, R^2^ = 0.04). The results of base pairing probability in the mRNAs having the top 1% enrichment factors, >1.00, <1.00, or the bottom 1% calculated by the ViennaRNA did not lead to a general conclusion (Fig. S3b).

### Method for comprehensive analysis of the effect of mRNA sequences on translation accuracy

To study the translation accuracy, researchers generally prepare a ribosome-paused state by combining ribosome, mRNA, aminoacyl-tRNAs, and related proteins^35^. In our translation system, we could design the ribosome-paused state at the UAG blank by avoiding the normal termination step involving RF-1^23,26^.

To study the effect of mRNA sequences on translation accuracy, we prepared mRNA libraries (fibronectin-UAG) carrying (VVN)_2_–UAG–Z (V represents a mixture of A, G, C; N represents a mixture of A, G, C, and U; Z represents N or no base; 6,480 sequences in theory) at the 3′-end (Fig. 3a and Data file S1). The first and second bases of codons were limited to V to avoid hydrophobic residues for further application as linkers to connect protein or peptide libraries to mRNAs. To avoid the undesired effect of amino acid sequences coded by VVN codons on pull-down efficiency, biotin was incorporated at the *N*-terminus of fibronectin using the Flexizyme system^36-38^, and only the fibronectin-PuL/mRNA complexes were isolated by the biotin pull-down^26,39^. An enrichment factor was calculated as described above. The enrichment factor mainly reflected the misreading efficiency of the UAG blank codon by near-cognate tRNAs because suppressing the UAG blank codon reduced the puromycin conjugation efficiency^26^. We also used an mRNA library (lipocalin-UAG) with a lipocalin-2 sequence to see whether the protein-coding sequence influences the effect of the two codons (VVN)_2_ and the single base Z on translation accuracy.

**Fig. 3.**
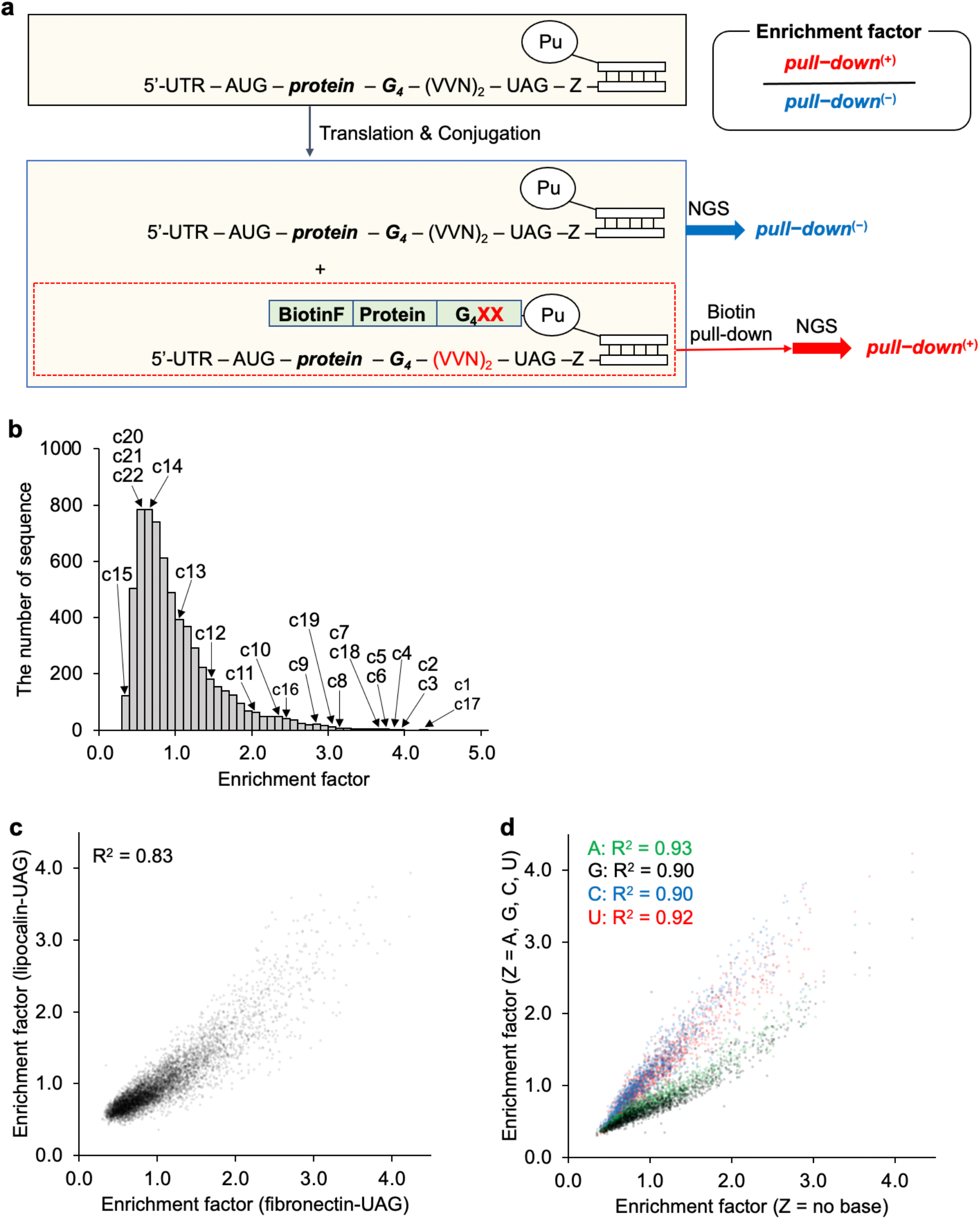
Comprehensive data sets of enrichment factors for mRNA libraries with VVN_2_-UAG-Z sequences carrying the different coding proteins to analyze the effect of the mRNA sequences on translation accuracy. **(a)** Method for analysis of the impact of mRNA sequences on translation elongation. Two codons and one base surrounding the UAG blank codon were randomized (VVN_2_-UAG-Z; V = A, C, G; N = A, U, G, C; Z = N or no base). Different protein-coding sequences (the fibronectin and the lipocalin) were used in each mRNA library. The PuL-attached mRNA library was added to the EF-2 depleted translation system, whose start codon was reprogrammed to biotin-Phe. Only the mRNAs with the coded protein were collected by pull-down using streptavidin. The enrichment factor was calculated from the ratio of the appearance rates before [*pull-down*^(−)^] and after the biotin pull-down [*pull-down*^(+)^]. **(b)** Histogram of enrichment factors of all the sequences in the fibronectin-UAG mRNA library. Enrichment factors of mRNAs used in Fig. 5 were labeled (c1–c22). **(c)** Scatter plot of enrichment factors from the fibronectin and lipocalin mRNA libraries, **(d)** mRNA carrying A, U, G, C, and no base after the UAG blank codon (Z position).

### The effect of RNA sequences on translation accuracy

We obtained more than 20 million reads before pull-down and after pull-down in both the fibronectin-UAG and lipocalin-UAG mRNA libraries (Data file S3). The enrichment factor ranged from 0.30 to 4.23, indicating the (VVN)_2_ codons had more impact on the enrichment factor than the three codons after the start codon (Fig. 3b vs. Fig. 1b and Fig. S4b vs. Fig. S1b). The enrichment factors were correlated well between the fibronectin-UAG and lipocalin-UAG mRNA libraries suggesting their coding sequence had less effect (Fig. 3c, R^2^ = 0.83). The base at the Z position after the UAG [(VVN)_2_–UAG–Z] had a significant impact on the enrichment factor (Figs. 3d and S4c). Transcripts with purine bases always produced lower enrichment factors than the ones with a pyrimidine base. The base after the amber codon reportedly affects the suppression efficiency of a cognate amber suppressor tRNA, and the purine base after the amber codon increases the suppression efficiency^40-44^ by stacking with the codon–anticodon duplex and stabilizing the codon–anticodon interaction^45^. In our case, misreading the UAG blank codon by near-cognate tRNAs^46,47^ competes with the puromycin reaction, causing a reduction in the enrichment factor. Our results suggested that the base after the UAG blank codon affected misreading rate in translation.

The heat maps of the enrichment factor on the (VVN)_2_ sequences suggested that the codons at the E-site and P-site independently affect the enrichment factor (Figs. 4, S5, and S6). mRNA having GGC and GGU codons at the E-site gave higher enrichment factors (Figs. S7a and S7b, upper panels), indicating that these tRNAs at the E-site tend to reduce misreading of the UAG blank codon at the A-site.

**Fig. 4.**
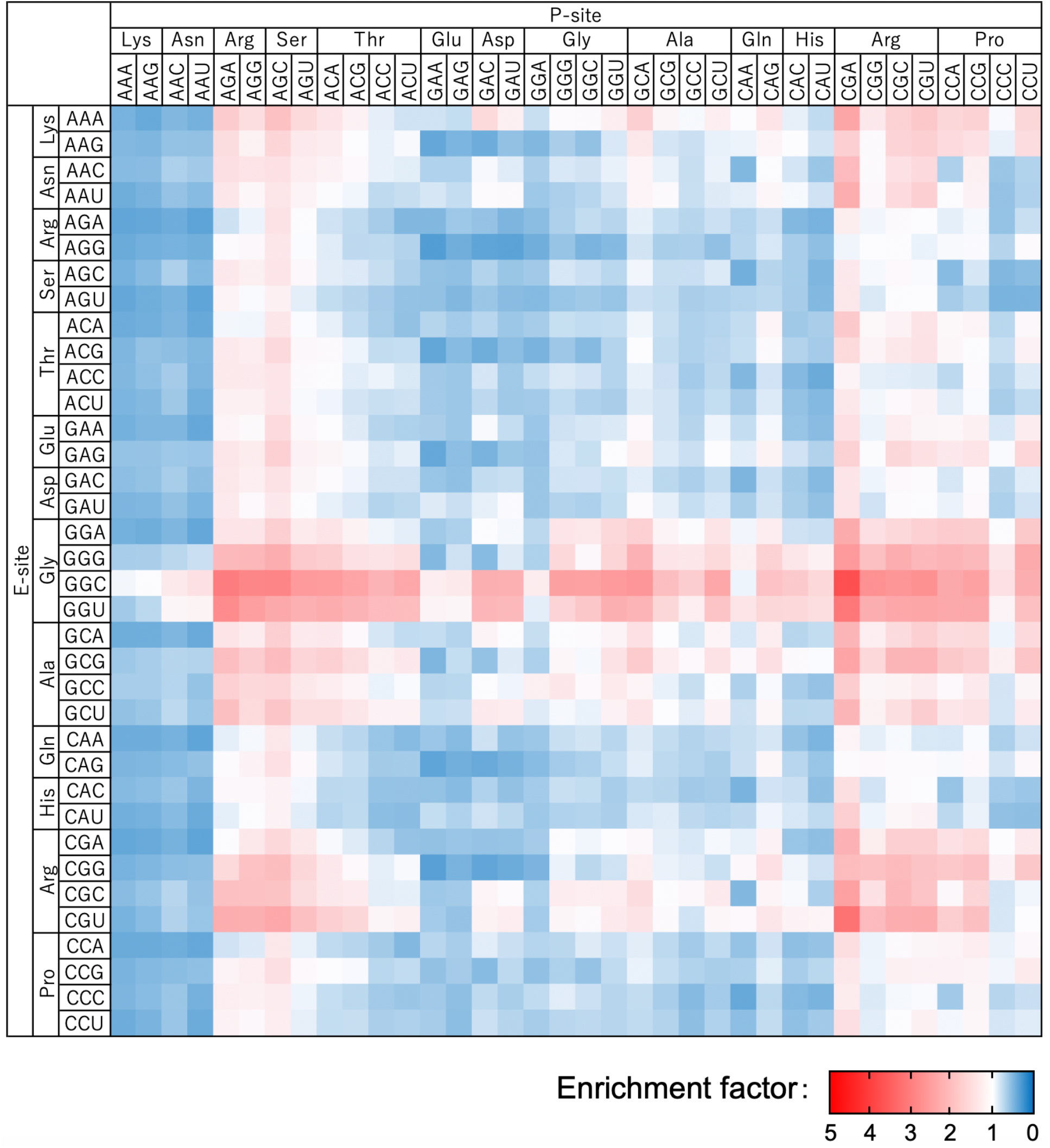
Heat map of enrichment factors of all fibronectin mRNAs carrying VVN_2_ codons before the UAG blank codon. The enrichment factors were the average of the ones from mRNA carrying all Z bases.

mRNAs having CGA at the P-site gave higher enrichment factors than others (Figs. S7a and S7b, lower panels). The CGA at the P-site, which forms I–A wobble pair causing the distorted anticodon loop^48^, inhibited the expression of proteins or reduced the elongation speed^49-51^. The CGA at the P-site would negatively affect the elongation reaction, thus reducing the misreading of the UAG blank codon by near-cognate tRNAs. mRNAs having AGC at the P-site also gave relatively higher enrichment factors than others. Those with AAA and AAG at the P-site gave significantly lower enrichment factors. These results might reflect the effect of these codons on the drop-off event induced by EF-G^52^. Since the *k*_off_ values of various peptidyl-tRNA are thought to be mostly uniform^53,54^, the impact of these codons at the P-site for the drop-off event needs to be studied further.

### Correlation between the enrichment factor from mRNA library and the puromycin conjugation efficiency of monotypic mRNA

To verify the correlation between the enrichment factor and the puromycin conjugation efficiency in monotypic mRNA, we selected 19 mRNAs (c1 to c19) that cover various enrichment factors (Fig. 3b and Table 1). We prepared each PuL/mRNA complex and added them to the cell-free translation system. After the reaction, the sample was loaded on Urea-SDS-PAGE. The puromycin conjugation efficiency was calculated from the amount of protein-PuL/mRNA complex over PuL and PuL/mRNA complex determined by analyzing the fluorescence emitted from the HEX-labeled PuL (Fig. 5a). The enrichment factor well correlated with the puromycin conjugation efficiency in monotypic mRNA (R^2^ = 0.83) (Fig. 5b).

**Table 1.**
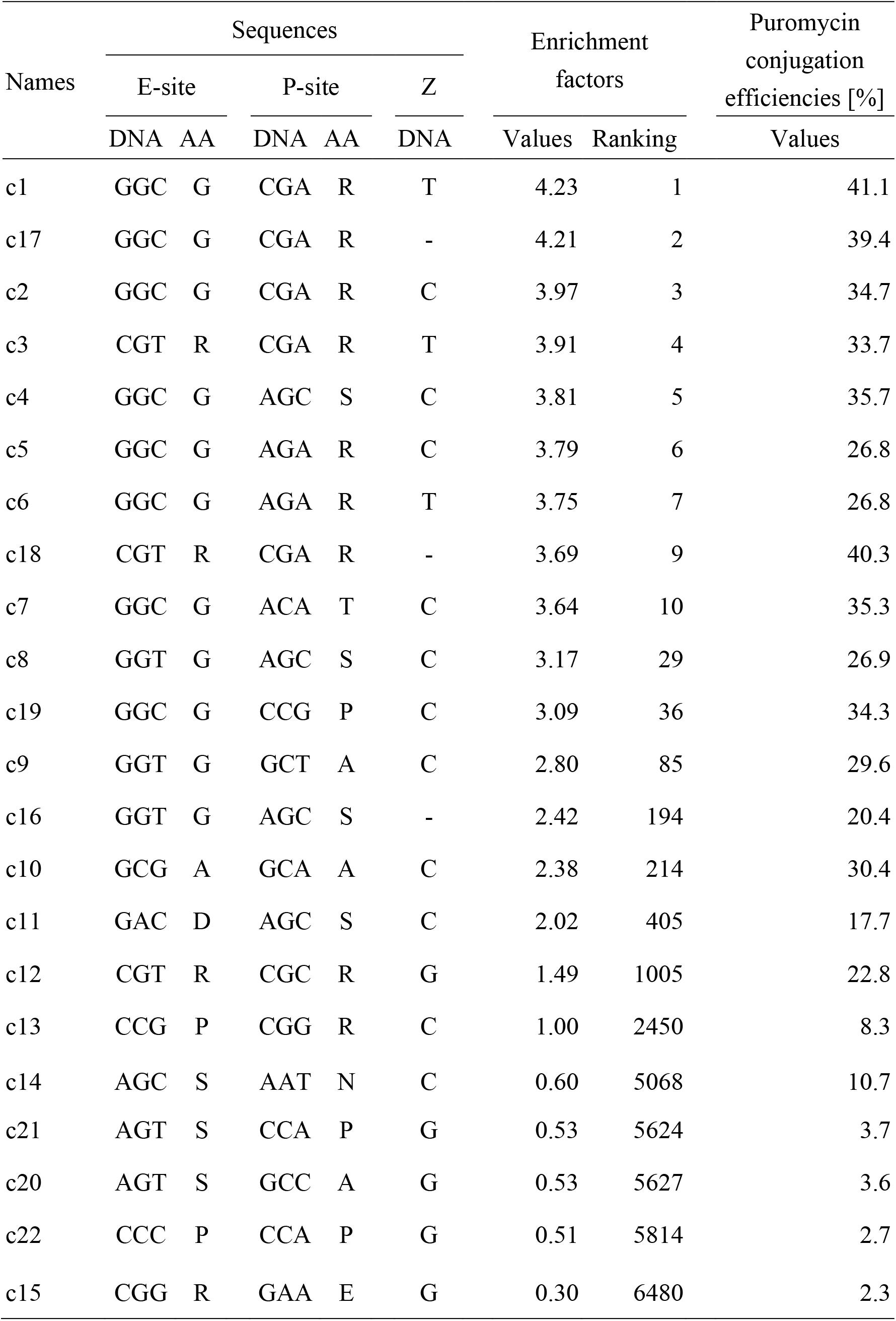
The 22 sequences around the UAG blank codon used to measure puromycin conjugation efficiency. The table contains the following information: sequences around the UAG blank codon, the values and rankings of enrichment factors in the SD8-fibronectin mRNA library, and the puromycin conjugation efficiencies of the SD8-fibronectin mRNA. Abbreviations: AA, amino acid.

| Names | Sequences |  |  |  |  | Enrichment factors |  | Puromycin conjugation efficiencies [%] |
| --- | --- | --- | --- | --- | --- | --- | --- | --- |
|  | E-site |  | P-site |  | Z |  |  |  |
|  | DNA | AA | DNA | AA | DNA | Values | Ranking | Values |
| c1 | GGC | G | CGA | R | T | 4.23 | 1 | 41.1 |
| c17 | GGC | G | CGA | R | - | 4.21 | 2 | 39.4 |
| c2 | GGC | G | CGA | R | C | 3.97 | 3 | 34.7 |
| c3 | CGT | R | CGA | R | T | 3.91 | 4 | 33.7 |
| c4 | GGC | G | AGC | S | C | 3.81 | 5 | 35.7 |
| c5 | GGC | G | AGA | R | C | 3.79 | 6 | 26.8 |
| c6 | GGC | G | AGA | R | T | 3.75 | 7 | 26.8 |
| c18 | CGT | R | CGA | R | - | 3.69 | 9 | 40.3 |
| c7 | GGC | G | ACA | T | C | 3.64 | 10 | 35.3 |
| c8 | GGT | G | AGC | S | C | 3.17 | 29 | 26.9 |
| c19 | GGC | G | CCG | P | C | 3.09 | 36 | 34.3 |
| c9 | GGT | G | GCT | A | C | 2.80 | 85 | 29.6 |
| c16 | GGT | G | AGC | S | - | 2.42 | 194 | 20.4 |
| c10 | GCG | A | GCA | A | C | 2.38 | 214 | 30.4 |
| c11 | GAC | D | AGC | S | C | 2.02 | 405 | 17.7 |
| c12 | CGT | R | CGC | R | G | 1.49 | 1005 | 22.8 |
| c13 | CCG | P | CGG | R | C | 1.00 | 2450 | 8.3 |
| c14 | AGC | S | AAT | N | C | 0.60 | 5068 | 10.7 |
| c21 | AGT | S | CCA | P | G | 0.53 | 5624 | 3.7 |
| c20 | AGT | S | GCC | A | G | 0.53 | 5627 | 3.6 |
| c22 | CCC | P | CCA | P | G | 0.51 | 5814 | 2.7 |
| c15 | CGG | R | GAA | E | G | 0.30 | 6480 | 2.3 |

**Fig. 5.**
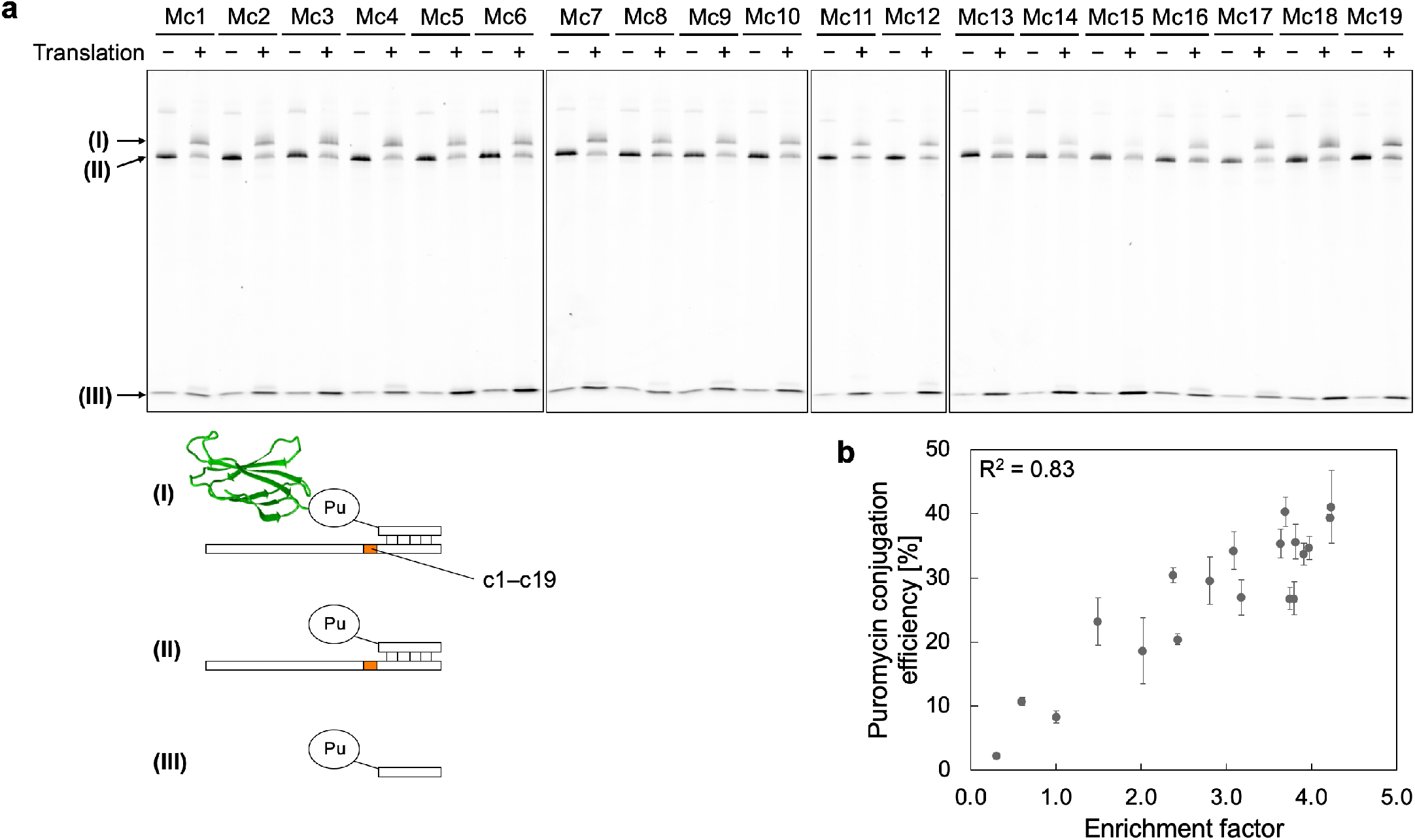
Puromycin conjugation efficiencies of various SD8-fibronectin mRNAs. **(a)** SDS-PAGE analysis of the puromycin conjugation efficiencies. The bands were detected using the fluorescence from HEX-labeled PuL. **(b)** Scatter plot between the enrichment factors and the puromycin conjugation efficiencies. Error bars indicate the ±SD of each experiment (*N* = 3).

### Effect of elongation factor P on puromycin conjugation efficiency

EF-P promotes peptide bond formation between Pro-tRNA^Pro^ and puromycin and consecutive proline residues by entropic steering of peptidyl-tRNA^Pro^ toward a catalytically productive orientation in the peptidyl transferase center of the ribosome^32^. We hypothesized that adding EF-P could accelerate the conjugation of PuL to proline at the end of the peptide, thus improving the puromycin conjugation efficiency. To test the hypothesis, we compared the puromycin conjugation efficiencies of two mRNAs carrying Arg (Mc1) and Pro (Mc19) in the presence or absence of EF-P. We found that the puromycin conjugation efficiencies with and without EF-P appeared to be the same (Fig. S8a). The puromycin conjugation efficiencies for Mc1 and Mc19 were relatively high (29%); therefore, we then used mRNAs (Mc20–c22) with lower puromycin conjugation efficiencies, but the efficiencies for each mRNA were similar with and without EF-P (Fig. S8b). The addition of EF-P might accelerate both the reaction of puromycin and the misreading of the UAG blank codon by near-cognate tRNAs.

### The identified sequences allowed the preparation of monobody and macrocyclic peptide libraries with high diversity

Puromycin conjugation efficiency is important in protein engineering because it determines the diversity of a protein library. To study the effect of sequences around the UAG blank on the preparation of a monobody library^23^, we prepared three mRNA libraries, each carrying one of the three sequences around the UAG blank codon that showed middle (1.00, c13, CCG-CGG-<u>UAG</u>-C, 2450th/6480), high (2.42, c16, GGU-AGC-<u>UAG</u>, 194th/6480), or very high (4.23, c1, GGC-CGA-<u>UAG</u>-U, 1st/6480) enrichment factors (Table 1). As a monobody library, we used BC-loop and FG-loop codon randomized genes (Fig. 6a). Higher puromycin conjugation efficiencies were observed for mRNA carrying c1 (MRc1, 20%) and c16 (MRc16, 18%) versus c13 (MRc13, 11%) (Fig. 6a). The puromycin conjugation efficiencies were partially correlated with the observations for fibronectin (11% vs. 8.3% for c13, 18% vs. 20.4% for c16, and 20% vs. 41.1% for c1; the monobody library vs. the fibronectin).

**Fig. 6.**
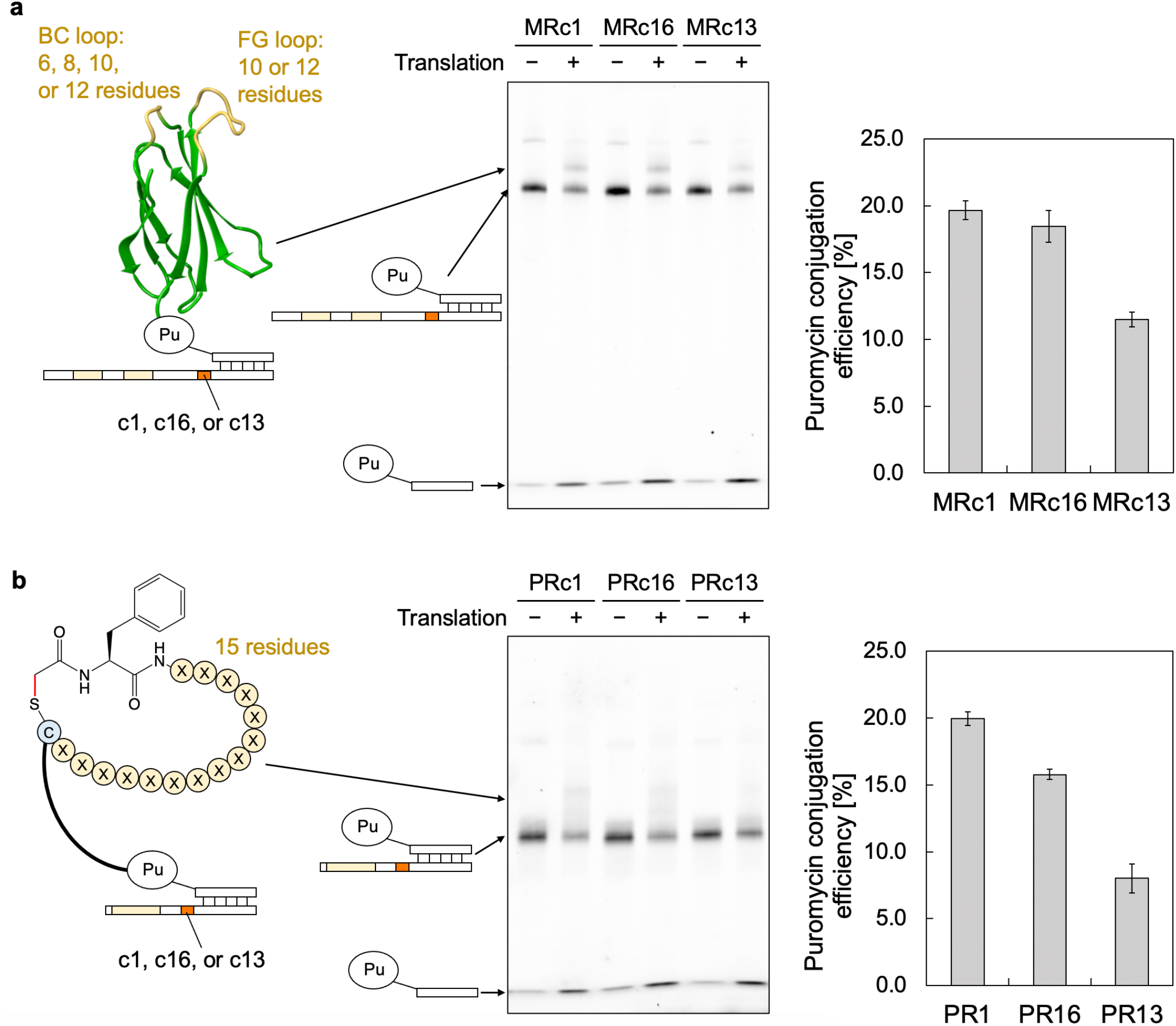
Puromycin conjugation efficiencies of the mRNA libraries. **(a)** Puromycin conjugation efficiencies of monobody libraries with randomized residues in the BC and FG loops (labeled in yellow) carrying c1, c16, or c13 sequences. **(b)** Puromycin conjugation efficiencies of macrocyclic peptide libraries with randomized 15 residues having c1, c16, or c13 sequences. The peptides were cyclized through a thioether bond formed by the reaction between the chloroacetyl group of *N*-terminal ClAcPhe and the thiol group of Cys. Error bars indicate the ±SD of each experiment (*N* = 3).

The effect of sequences around the UAG blank on the enrichment factor did not vary with protein-coding sequence (Fig. 3c). This fact led us to study the effect of sequences around the UAG blank on preparing a macrocyclic peptide library^26,55-58^. We prepared thioether cyclized macrocyclic peptide libraries^59^ (Fig. 6b) using the Flexizyme system^36-38^. Higher puromycin conjugation efficiencies were observed for mRNA carrying c1 (PRc1, 20%) and c16 (PRc16, 16%) versus c13 (PRc13, 8%) (Fig. 6b).

In both monobody and macrocyclic peptide libraries, the sequences around the UAG blank codon with high (2.42, c16, 194th/6480) and very high (4.23, c1, 1st/6480) enrichment factors gave 16% to 20% puromycin conjugation efficiencies. We think at least the top 3% of sequences (194 sequences) could be useful for preparing highly diverse libraries of various scaffold proteins and macrocyclic peptides in future.

## Discussion

In previous protein engineering studies, *N*-terminal and *C*-terminal sequences (7–12 residues in total) were selected *in vitro* via mRNA display to increase puromycin conjugation efficiency^60,61^. Those sequences increased puromycin conjugation efficiencies up to 40% for a single-chain antibody^60^ and 23% for a variable domain of heavy chain antibody (VHH)^61^. By contrast, the selected sequence for a VHH did not improve other VHH variants and other backbone proteins. A similar mRNA display method using the Flexizyme system was recently applied to investigate the effect of one to three residues before the Pro_3_ sequence on the drop-off– reinitiation event^62^.

In this study, we used mRNA display to conduct a comprehensive analysis of the effect of mRNA sequences on translation efficiency and accuracy. We first prepared three mRNA libraries (SD8-fibronectin, SD8-lipocalin, and SDcat-fibronectin; 32,768 × 3 sequences in theory) carrying three NNW codons after the start codon. The comparison of NGS sequences between the translated mRNAs and the original mRNA libraries revealed that the effect of each three-codon sequence on the enrichment factor was strongly dependent on the protein-coding sequence (Fig. 1d, R^2^ = 0.08) as well as the 5′-UTR sequence (Fig. 1e, R^2^ = 0.36). The data analysis, including the base content, the codon content, the amino acid contents, the calculated folding free energy of the mRNA structure, and the mRNA base pair probability, gave no determinate factor to explain the relationship between each three-codon sequence and the enrichment factor. This contradicts previous studies that used FACS^9-13^ or multiplex automated genome engineering mutagenesis with amplicon deep-sequencing^63^, which suggested a weak to middle correlation between the calculated folding free energy of the mRNA structure and the translation efficiency. In this study, the effect of the folding of mRNA on the translation efficiency might be minimized by at least the following two reasons; (1) weaker mRNA secondary structures with A or U bases at the third base of codons than that with G or C bases, (2) no involvement of other factors, such as ribonucleases and proteases, which cause mRNA and protein degradation.

For study of translation accuracy, we prepared two mRNA libraries (the fibronectin-UAG and the lipocalin-UAG) carrying (VVN)_2_–UAG–Z (6,480 × 2 sequences in theory) at the 3′-end region. The comparison of NGS sequences between the translated mRNAs and the original mRNA libraries showed that the effect of each sequence on the enrichment factor was independent of the protein-coding sequence (Fig. 3c, R^2^ = 0.83). The data analysis revealed that the purine base after the UAG blank codon increased decoding errors in the translation (Figs. 3d, S4c). The E-site and P-site codon independently influenced their enrichment factors (Fig. 4, S5, and S6). The GGC and GGU codons at the E-site gave higher enrichment factors than others (Figs. S7a and S7b, upper panels). As we described above, the presence of tRNA at the E-site is known to induce a lower affinity binding state of the A-site. This result might suggest that tRNAs corresponding to these codons have a higher affinity at the E-site than other tRNAs and prevent the misreading of the UAG blank codon by near-cognate tRNAs. The CGA at the P-site gave higher enrichment factors than others (Figs. S7a and S7b, lower panels). We think that the CGA at the P-site affected the elongation reaction negatively, thus reducing the misreading of the UAG blank codon by near-cognate tRNAs.

The study of the translation accuracy taught us *C*-terminal sequences that had high enrichment factors for fibronectin and lipocalin coding mRNAs. We demonstrated the construction of highly-diverse monobody and macrocyclic peptide libraries via the obtained *C*-terminal sequences. These libraries will facilitate the further development of functional molecules that will contribute to biological research and therapeutic applications.

## Methods

### Materials

The oligonucleotides and synthetic DNA templates were purchased either Fasmac Co., Ltd. (Japan), Nippon Bio Service (Japan) or Integrated DNA Technologies (IA, USA). The sequences were listed in Data file S4. The restriction enzymes were obtained from New England Biolabs (MA, USA). The preparation of the cell-free translation system, *Pfu-S* DNA polymerase, and Moloney murine leukemia virus reverse transcriptase (MMLV) were described in a previous report^19-23^.

### Preparation of mRNA libraries with random sequence following start codon

To prepare DNA libraries of SD8-fibronectin, 3.5 fmol of MonoS(H)SSS-HA gene was added to the reaction mixture A [10 mM Tris-HCl pH 8.4, 100 mM KCl, 0.1% (v/v) Triton X-100, 2% (v/v) DMSO, 2 mM MgSO_4_, 0.2 mM each dNTP, and 2 nM of *Pfu-S* DNA polymerase] containing 0.375 μM MonoS(H)SSS.F26 and 0.375 μM HATag-G4S.R48, and amplified by PCR (18 μL in total, ten cycles of PCR). The mixture (0.16 μL) was added to the reaction mixture A containing 0.375 μM MonoS(H)NNW3SSS.F58 and 0.375 μM G5S-4Gan21-3.R42, and amplified by PCR (25 μL in total, nine cycles of PCR). The mixture (0.16 μL) was added to the reaction mixture A containing 0.375 μM T7SD8M2.F44 and 0.375 μM G5S-4Gan21-3.R42, and amplified by PCR (160 μL in total, ten cycles of PCR). The amplified DNA library was purified by phenol/chloroform extraction and isopropanol precipitation.

The DNA libraries of SDcat-fibronectin was prepared as similar protocol as above except the primers (catMonoS(H)NNW3SSS.F60 instead of MonoS(H)NNW3SSS.F58 and T7SDCATM.F46 instead of T7SD8M2.F44 were used).

The DNA libraries of SD8-lipocalin was prepared as similar protocol as above except the gene (Ant-wt-speI gene instead of MonoS(H)SSS-HA gene) and the primers (Lcn.F22 instead of MonoS(H)SSS.F26, Lcn-HATag.R73 instead of HATag-G4S.R48, LcnNNW3.F54 instead of MonoS(H)NNW3SSS.F58, and HATag-G4S.R48 instead of G5S-4Gan21-3.R42)

These DNA templates were transcribed by *in vitro* run-off transcription under the following conditions: 40 mM Tris-HCl pH 8.0, 1 mM spermidine, 0.01% (v/v) Triton X-100, 10 mM DTT, 30 mM MgCl2, 5 mM of each NTP, the DNA library, and 0.18 μM T7 RNA polymerase. The mRNA was purified by phenol/chloroform extraction and isopropanol precipitation. The mRNA concentration was determined by OD at 260 nm.

### Preparation of mRNA libraries with random sequence around UAG blank codon

To prepare DNA libraries of fibronectin-UAG, 3.5 fmol of MonoS(H)SSS-HA gene was added to the reaction mixture A containing 0.375 μM SD8-MQANSGS-MonoS(H).F61 and 0.375 μM MonoS(H)-GGG.R32, and amplified by PCR (18 μL in total, ten cycles of PCR). The mixture (0.16 μL) was added to the reaction mixture A containing 0.375 μM T7SD8M2.F44, 0.069 μM MonoS(H)-VVN2.R48, and 0.276 μM MonoS(H)-VVN2N1.R49, and amplified by PCR (160 μL in total, nine cycles of PCR). The amplified DNA library was purified by phenol/chloroform extraction and isopropanol precipitation.

The DNA libraries of lipocalin-UAG was prepared as similar protocol as above except the gene (Ant-wt gene instead of MonoS(H)SSS-HA gene) and the primers (SD8-Lcn.F62 instead of SD8-MQANSGS-MonoS(H).F61, Lcn-GGG.R33 instead of MonoS(H)-GGG.R32, Lcn-VVN2.R48instead of MonoS(H)-VVN2.R48, and Lcn-VVN2N1.R49 instead of MonoS(H)-VVN2N1.R49)

These DNA templates were transcribed as the same protocol described as above.

### Analysis of enrichment factors for mRNA libraries with random sequence following start codon

The PuL/mRNA complex was prepared by annealing of Hex-Pu-an21-3 (4 μM) and mRNA (6.7 μM) in an annealing buffer (25 mM HEPES-K pH 7.8, 200 mM potassium acetate) by heating the solution (5 μL) to 95°C for 2 min; it was the cooled down to 25 °C. The resulting solution (1 μL) was added to a reconstituted translation system without RF-1. The reaction mixture (4 μL) was incubated at 37°C for 30 min. After the reaction, 0.8 μL of 100 mM EDTA (pH 8.0) was added to the translation mixture. The resulting solution was diluted with reverse transcription mixture A [2.4 μL; 150 mM Tris-HCl pH 8.4, 225 mM KCl, 75 mM MgCl_2_, 16.5 mM dithiothreitol (DTT), 1.5 mM dNTPs, 0.23 μM MMLV, 7.5 μM G4S.R19], and the resulting solution was incubated at 42°C for 15 min. The buffer was exchanged to HBST buffer [50 mM Hepes-KOH (pH 7.5), 300 mM NaCl, 0.05% (v/v) Tween 20] using Zeba Spin Desalting Columns. The entire solution was mixed with 1 μL of Anti-HA tag mAb (TANA2) (Medical & Biological Laboratories Co., Ltd) and the resulting solution was incubated at 25°C for 30 min. In order to pull down protein-PuL/mRNA complexes, the resulting solution were mixed with 20 μL of Dynabeads Protein G for immunoprecipitation (Thermo Fisher Scientific) at 25°C for 10 min. The pull-down beads were washed with 10 μL of HBST buffer for 10 s.

The solution before the pull-down was diluted 100 times with 1xPCR dNTPs [10 mM Tris-HCl pH 8.4, 100 mM KCl, 0.1% (v/v) Triton X-100, 2 mM MgSO4, 0.22 mM dNTPs]. All the beads after the pull-down were added to 100 μL of 1xPCR dNTPs, followed by being heated at 95 °C for 5 min. The diluted solution before the pull-down and only supernatant of the mixture after the pull-down (85 μL each) were added to PCR premix [10 mM Tris-HCl pH 8.4, 100 mM KCl, 0.1% (v/v) Triton X-100, 4% (v/v) DMSO, 2 mM MgSO_4_, 0.2 Mm each dNTP, and 4 nM of *Pfu-S* DNA polymerase] (85 μL), PCR-amplified using primer mix (Table S1), and purified by phenol/chloroform extraction and isopropanol precipitation. The sequences of the DNAs were analyzed using the NovaSeq6000 platform (Illumina) at Macrogen (Japan). Appearance rates of sequences were calculated as a percentage of the read counts of the sequences in the total read counts in the whole library. The appearance rate of each sequence after pull-down was divided by the corresponding one before pull-down to determine the enrichment.

### Analysis of enrichment factors for mRNA libraries with random sequence around UAG blank codon

The PuL/mRNA complex was prepared by annealing of Hex-Pu-an21-3 (4 μM) and mRNA (6.7 μM) in an annealing buffer by heating the solution (5 μL) to 95°C for 2 min; it was the cooled down to 25 °C. The resulting solution (1 μL) was added to a reconstituted translation system that contained 16 μM biotin-phenylalanine-tRNA^fMet^_CAU_ and did not contain RF1 and formyl donor (FD). The reaction mixture (4 μL) was incubated at 37°C for 30 min. After the reaction, 1 μL of 100 mM EDTA (pH 8.0) was added to the translation mixture. The resulting solution was diluted 50 times with HBST buffer. In order to pull down protein-PuL/mRNA complexes, 10 μL of the diluted solution was mixed with 20 μL of Dynabeads M-280 streptavidin (Thermo Fisher Scientific) at 25°C for 5 min. The pull-down beads were washed with 50 μL of HBST buffer for 1 min, and 50 μL of HBS buffer [50 mM Hepes-KOH (pH 7.5), 300 mM NaCl] for 1 min.

The solution before the pull-down was diluted 50 times with reverse transcription mixture B [150 mM Tris-HCl pH 8.4, 225 mM KCl, 75 mM MgCl_2_, 16.5 mM dithiothreitol (DTT), 1.5 mM dNTPs, 0.23 μM MMLV] The diluted solution before the pull-down (1 μL) or all the beads after the pull-down were added to an annealing buffer containing 5 μM an21-3.R21 (10 μL each in total), followed by being heated at 95 °C for 2 min. After 30 s incubation at 25 °C, the mixture before the pull-down and only supernatant of the mixture after the pull-down were diluted with reverse transcription mixture C [10 μL; 124 mM Tris-HCl pH 8.4, 186 mM KCl, 62.2 mM MgCl_2_ and 13.7 mM dithiothreitol (DTT), 1.24 mM dNTPs, and 0.23 μM MMLV], and the resulting solution was incubated at 42°C for 30 min.

These solutions were diluted with 180 μL of 1xPCR dNTPs, and 85 μL of them were added to PCR premix (85 μL), PCR-amplified using primer mix (Table S1), and purified by phenol/chloroform extraction and isopropanol precipitation. The protocols for sequencing the DNAs and calculating enrichment factors were same as above.

### Preparation of mRNAs for analysis of puromycin-conjugation efficiencies

To prepare DNAs of Mc1-Mc22, 7.5 fmol of MonoS(H)WTproduct gene was added to the reaction mixture A containing 0.375 μM MoS-QANSGS.F62 and 0.375 μM MonoS(H)-GGG.R32, and amplified by PCR (20 μL in total, 10 cycles of PCR). The mixture (0.16 μL) was added to the reaction mixture A containing 0.375 μM T7SD8M2.F44 and 0.375 μM reverse primer M (Table S2), and amplified by PCR (160 μL in total, 12 cycles of PCR). The amplified DNA was purified by phenol/chloroform extraction and isopropanol precipitation.

The DNA libraries of MRc1, MRc13, and MRc16 were prepared as similar protocol as above except the following conditions: MoS-pool-1 gene was used instead of MonoS(H)WTproduct gene; the reverse primer MR (Table S3) were used instead of reverse primer M; and numbers of cycles of 2nd PCR was decreased from 12 cycles to 10 cycles.

To prepare DNA libraries of PRc1, PRc13, and PRc16, 7.5 fmol of Peptide-pool(n=15) gene was added to the reaction mixture A containing 0.375 μM T7SD8M2.F44 and 0.375 μM reverse primer PR (Table S4), and amplified by PCR (160 μL in total, 10 cycles of PCR). The amplified DNA was purified by phenol/chloroform extraction and isopropanol precipitation.

The DNAs and DNA libraries were transcribed into mRNA as the same protocol as above.

### Measurement of puromycin conjugation efficiency

The PuL/mRNA complex was prepared by annealing PuL (4 μM) and mRNA (4.8 μM) in annealing buffer (25 mM HEPES-K pH 7.8, 200 mM potassium acetate) by heating the solution (4.8 μL) to 95°C for 2 min and cooling to 25°C. The resulting complex (1 μM) was added to a reconstituted translation system that RF1 was excluded. The reaction mixture (5 μL) was incubated at 37°C for 30 min. The reaction mixture (1 μL) was collected before and after translation and were mixed with 11 μL of gel shift buffer [62.5 mM Tris-HCl pH 6.8, 5 mM DTT, 0.05% (w/v) SDS, 10 mM MgCl_2_, 20% (v/v) Glycerol]. The samples (2 μL) were loaded onto 8% PAGE containing 0.375 M Tris-HCl pH 8.8, 0.1 % (w/v) SDS, and 6 M urea. The resulting gel was analyzed using the ChemiDoc MP Imaging System (Bio-Rad). The puromycin conjugation efficiency was calculated from the amount of protein-PuL/mRNA complex over PuL and PuL/mRNA complex determined by analyzing the fluorescence emitted from the HEX-labeled PuL.

For other puromycin conjugation study, following conditions were modified: For measurement of puromycin conjugation efficiency in the cell-free translation system with EF-P, 3 μM EF-P was added to the reconstituted translation system. For monobody library, the incubation time was changed from 30 min to 10 min. For macrocyclic peptide library, 10 μM *N*-chloroacetyl-L-phenylalanine-tRNA^fMet^_CAU_ was added to the reconstituted translation system. In addition, formyl donor was excluded from the translation mixture. 8% PAGE containing 0.45 M Tris-HCl pH 8.8, 0.1 % (w/v) SDS, and 6 M urea was used for electrophoresis for the macrocyclic peptide libraries.

### Calculation of minimum free energy of RNA secondary structure

Minimum free energies of RNA secondary structure (5’-UTR followed by 112 bases of the protein coding region) and base pairing probability were calculated using the RNAfold program from the ViennaRNA package^34^. The used python codes were listed in Data file S5 and S6.

## Acknowledgments

We thank Ms. Minori Eguchi and Mr. Seita Kito for their contributions to the early stages of this work. This study was financially supported by AMED (grant numbers 20he0622010h0001 and 20fk0108293s0101 to H.M.), a Grant-in-Aid for Scientific Research on Innovative Areas (grant number 20H04704 to G.H.), and a Grant-in-Aid for Early-Career Scientists (grant number 18K14332 to T.F.) from the Japan Society for the Promotion of Science, and a donation from H. Murakami.

## Contributions

S.U. carried out most of the experiments. T.K. did the early experiments. S.U. and H.M. wrote the manuscript with support from T.K. and T.F., and G.H. H.M. supervised the project.

## Additional information

All data needed to evaluate the conclusions in the paper are present in the paper and the Supplementary Materials. Requests for the genes should be submitted to H.M.

## Supplementary information

Data_file_bioRxiv.xlsx

## Figures and Tables

**Fig. S1.**
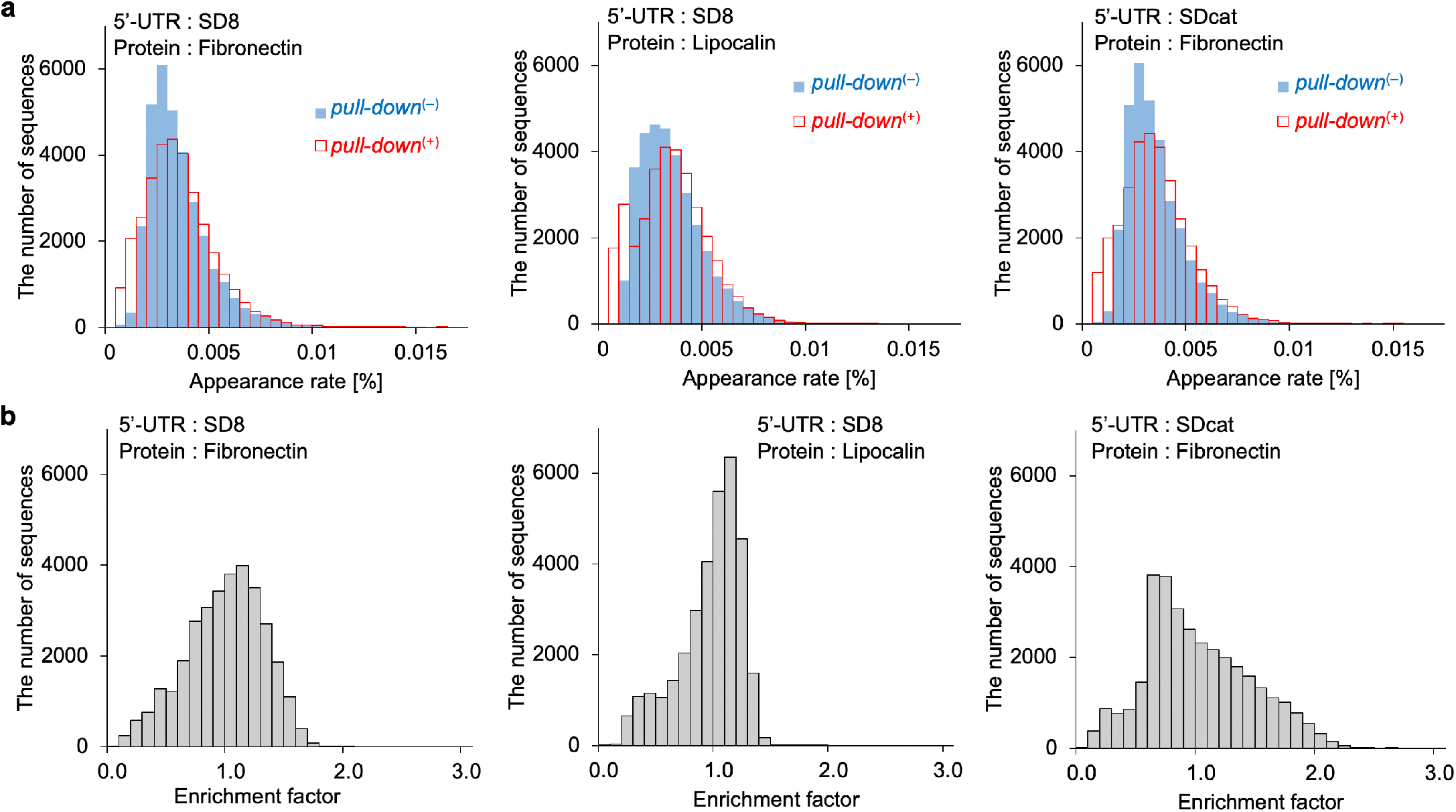
Data sets of enrichment factors for mRNA libraries with AUG-NNW_3_ codons carrying the different 5′-UTRs and protein coding sequences to analyze the effect of the mRNA sequences on translation efficiency. **(a)** Histogram of appearance rates of mRNAs before and after pull-down. **(b)** Histogram of enrichment factors of mRNAs. Three mRNA libraries (the SD8-fibronectin, the SD8-lipocalin, and the SDcat-fibronectin) were used. The left panel of (b) was also shown in Fig. 1.

**Fig. S2.**
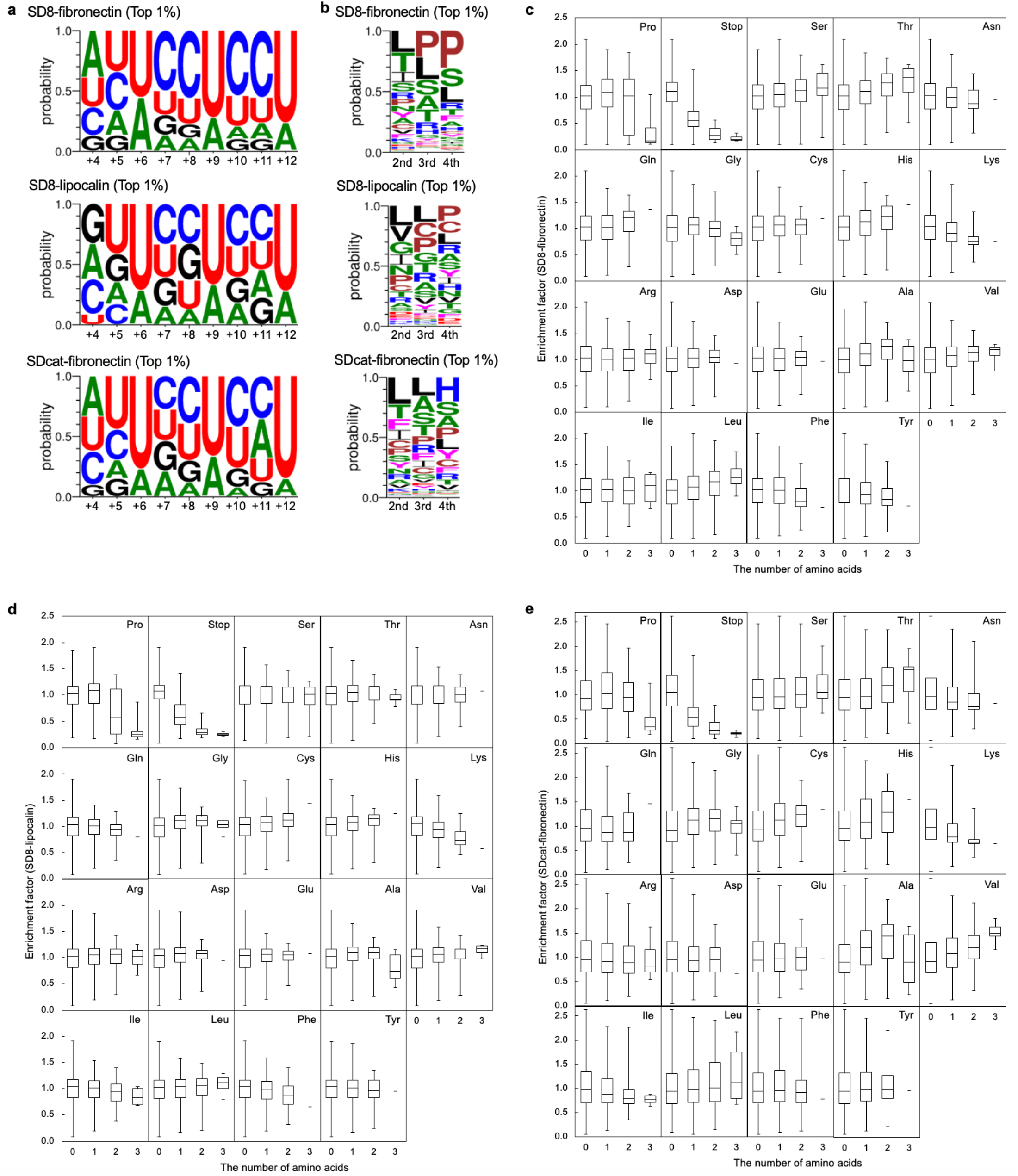
Comprehensive analysis of mRNA sequences on translation efficiency. Appearance frequencies of **(a)** nucleotides and **(b)** residues at each randomized position in the mRNA having top 1% enrichment factors (327 sequences). Box plots showing distributions of enrichment factors categorized according to the number of each amino acid in the random region following the start codon in **(c)** the SD8-fibronectin, **(d)** the SD8-lipocalin, and **(e)** the SDcat-fibronectin mRNA libraries.

**Fig. S3.**
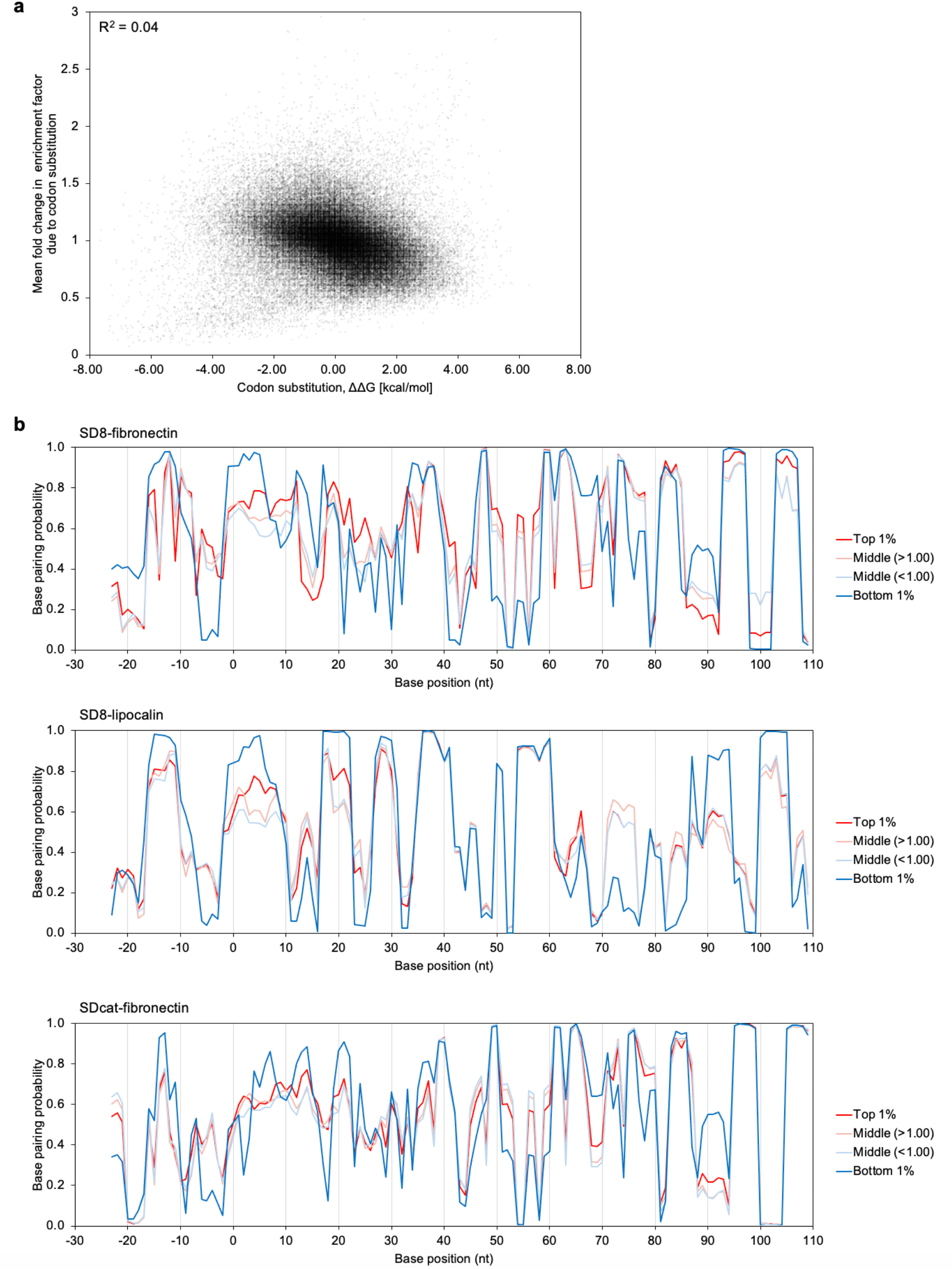
Analysis of mRNA folding. **(a)** The folding energy difference of synonymous codon variants and the mean fold change in enrichment factors. **(b)** Base pairing probability in the mRNAs having top 1% enrichment efficiencies, more than 1.00 enrichment efficiency, less than 1.00 enrichment efficiency, and bottom 1% enrichment efficiencies. Calculations were performed by the ViennaRNA^34^.

**Fig. S4.**
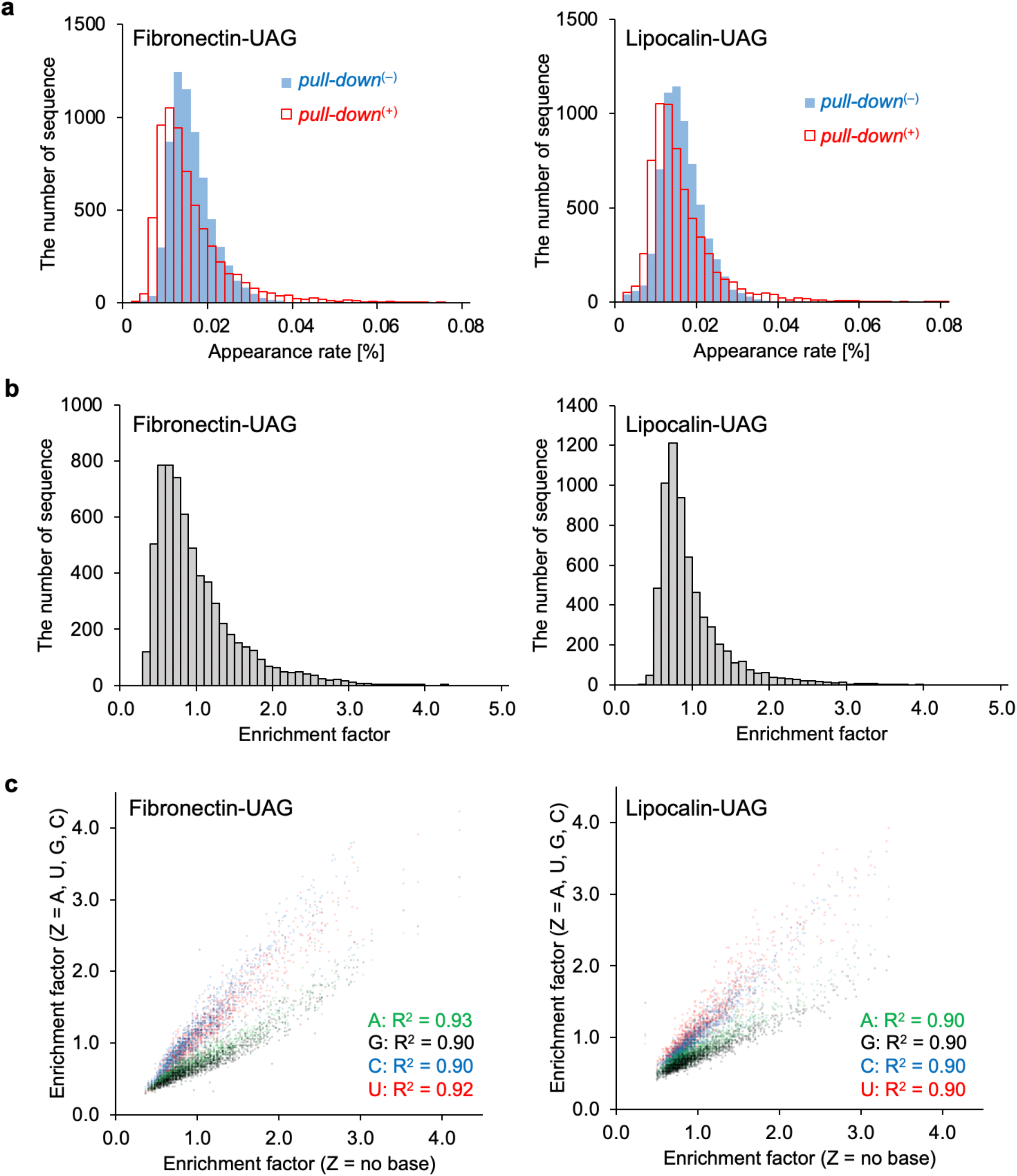
Comprehensive data sets of enrichment factors for mRNA libraries with VVN_2_-UAG-Z sequences carrying the different coding proteins to analyze the effect of the mRNA sequences on translation accuracy. **(a)** Histogram of appearance rates of mRNAs before and after pull-down. **(b)** Histogram of enrichment factors of mRNAs. **(c)** Scatter plot of enrichment factors of mRNA carrying A, U, G, C, and no base after the UAG blank codon (Z position). Two mRNA libraries (the fibronectin-UAG and the lipocalin-UAG) were used. Left panels of (b) and (c) were also shown in Fig. 3.

**Fig. S5.**
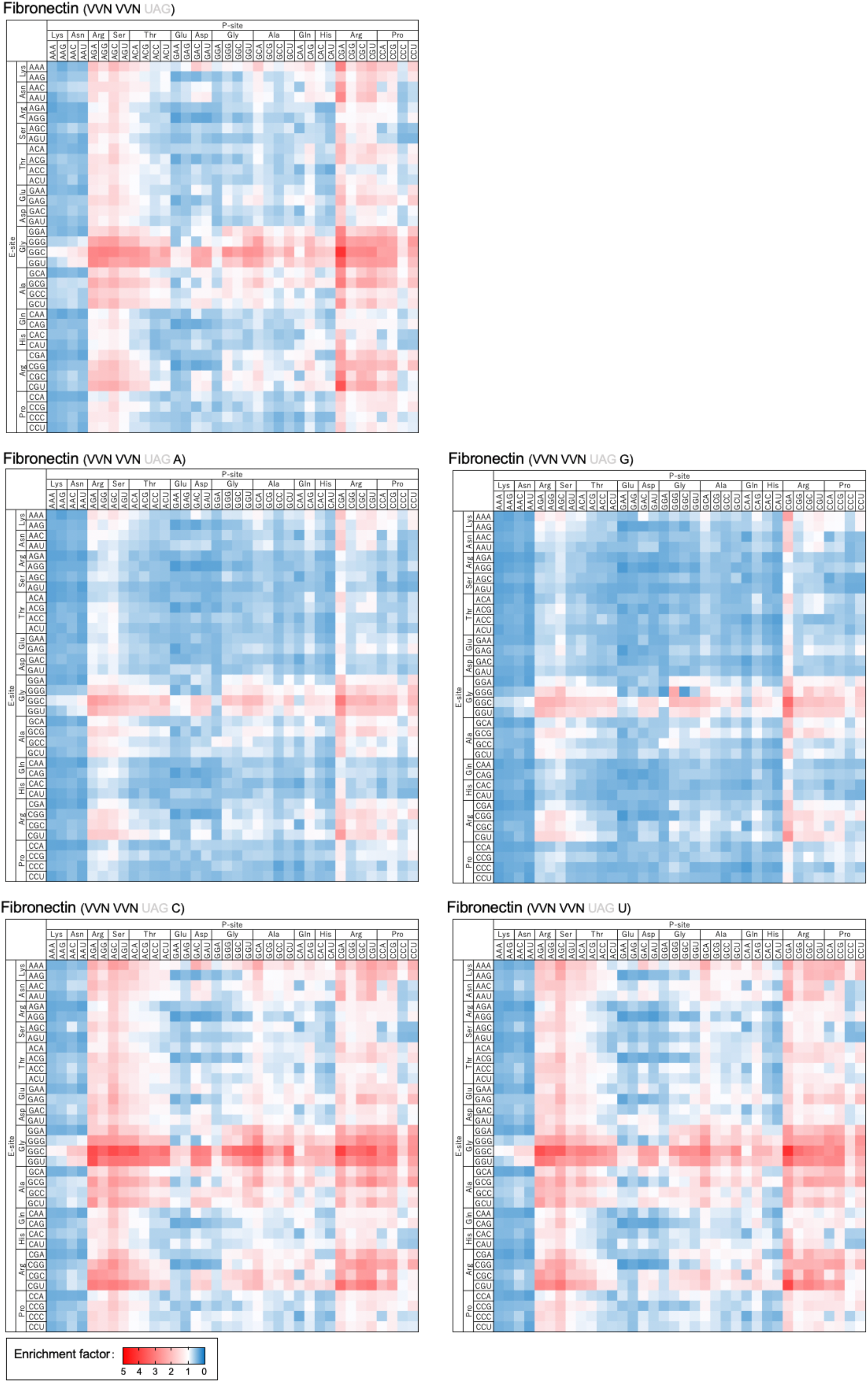
Heat maps of enrichment factors of all fibronectin mRNAs carrying VVN_2_ codons before the UAG blank codon. Results from mRNAs carrying A, U, G, C, and no base at the Z position were separately shown.

**Fig. S6.**
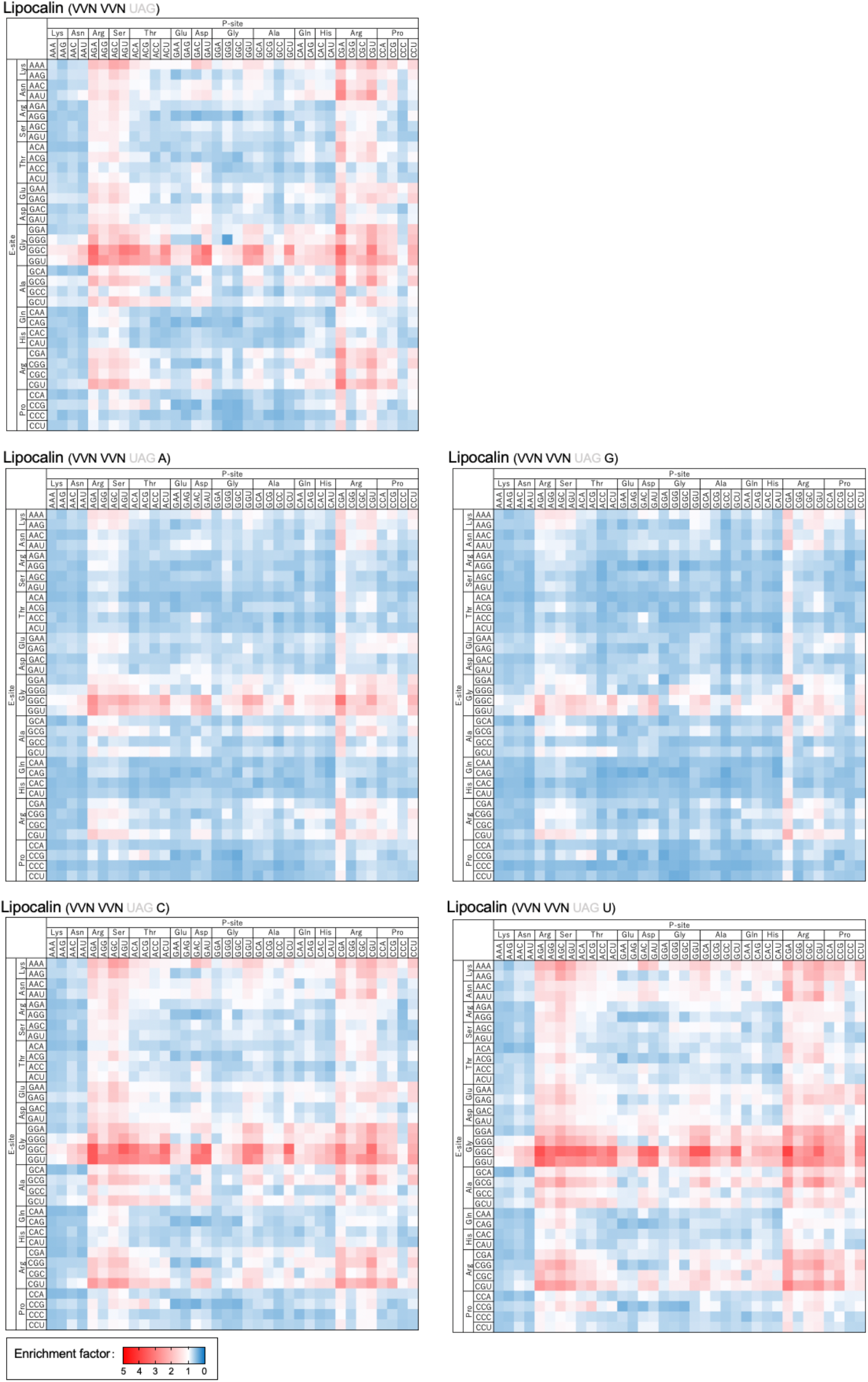
Heat maps of enrichment factors of all lipocalin mRNAs carrying VVN_2_ codons before the UAG blank codon. Results from mRNAs carrying A, U, G, C, and no base at the Z position were separately shown.

**Fig. S7.**
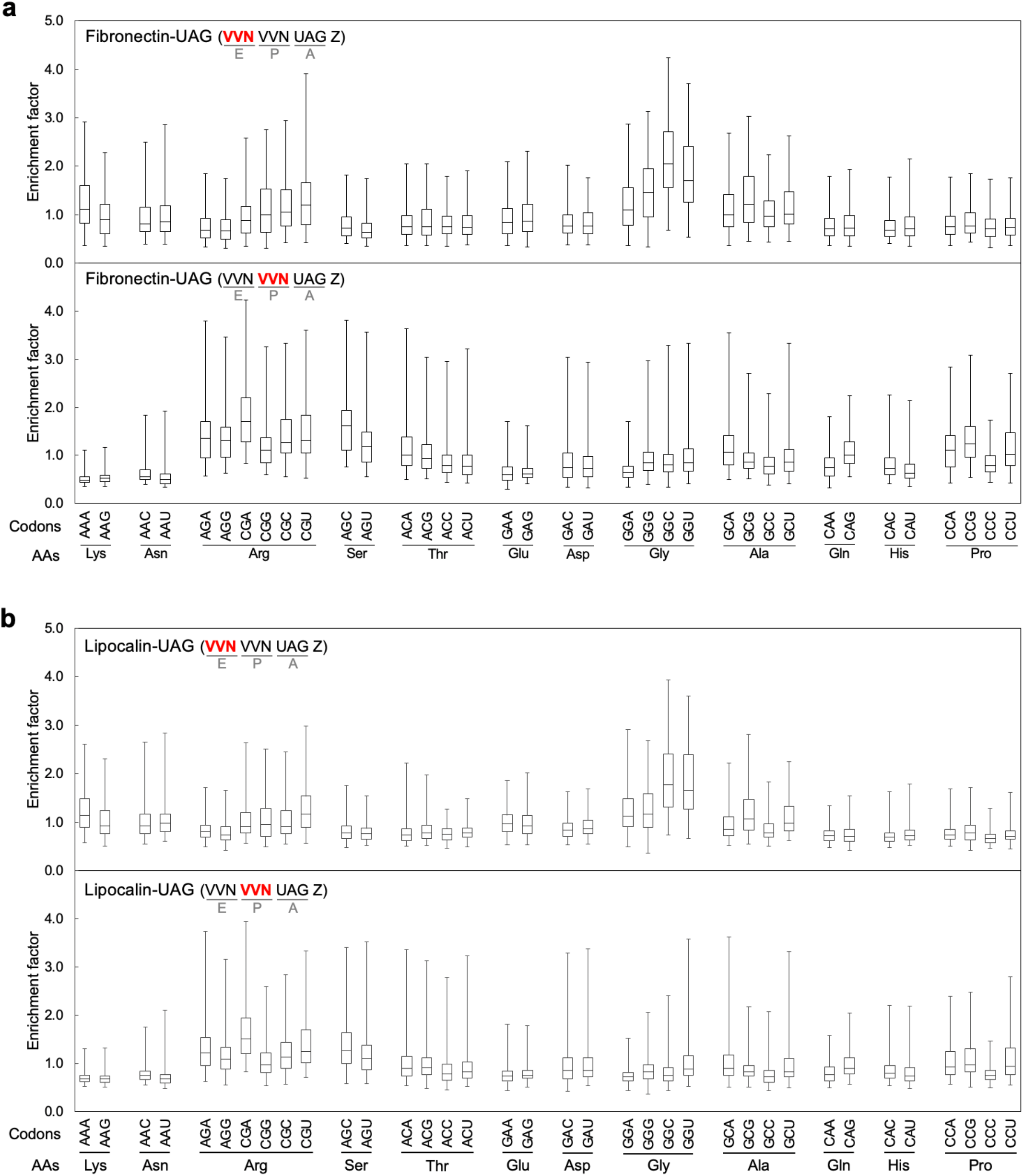
Box plots show distributions of enrichment factors categorized according to the codon at each position. **(a)** the fibronectin-UAG mRNA library. **(b)** the lipocalin-UAG mRNA library.

**Fig. S8.**
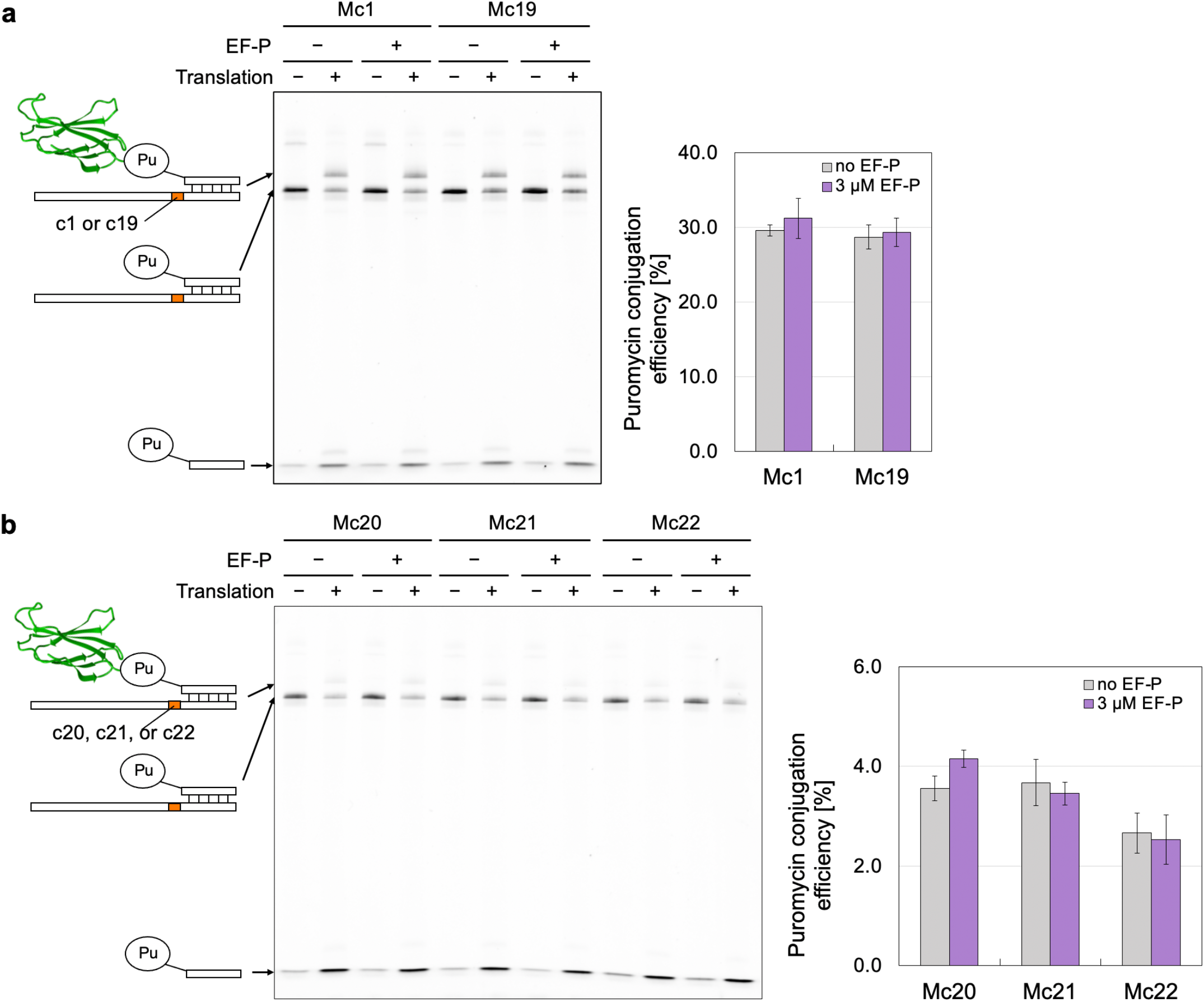
Effect of EF-P on the puromycin conjugation efficiencies. **(a)** Mc1 (no proline) and Mc19 (with one proline at P-site). **(b)** Mc20 (no proline), Mc21 (one proline at P-site), and Mc22 (two prolines). Error bars indicate the ±SD of each experiment (*N* = 3).

**Fig. S9.**
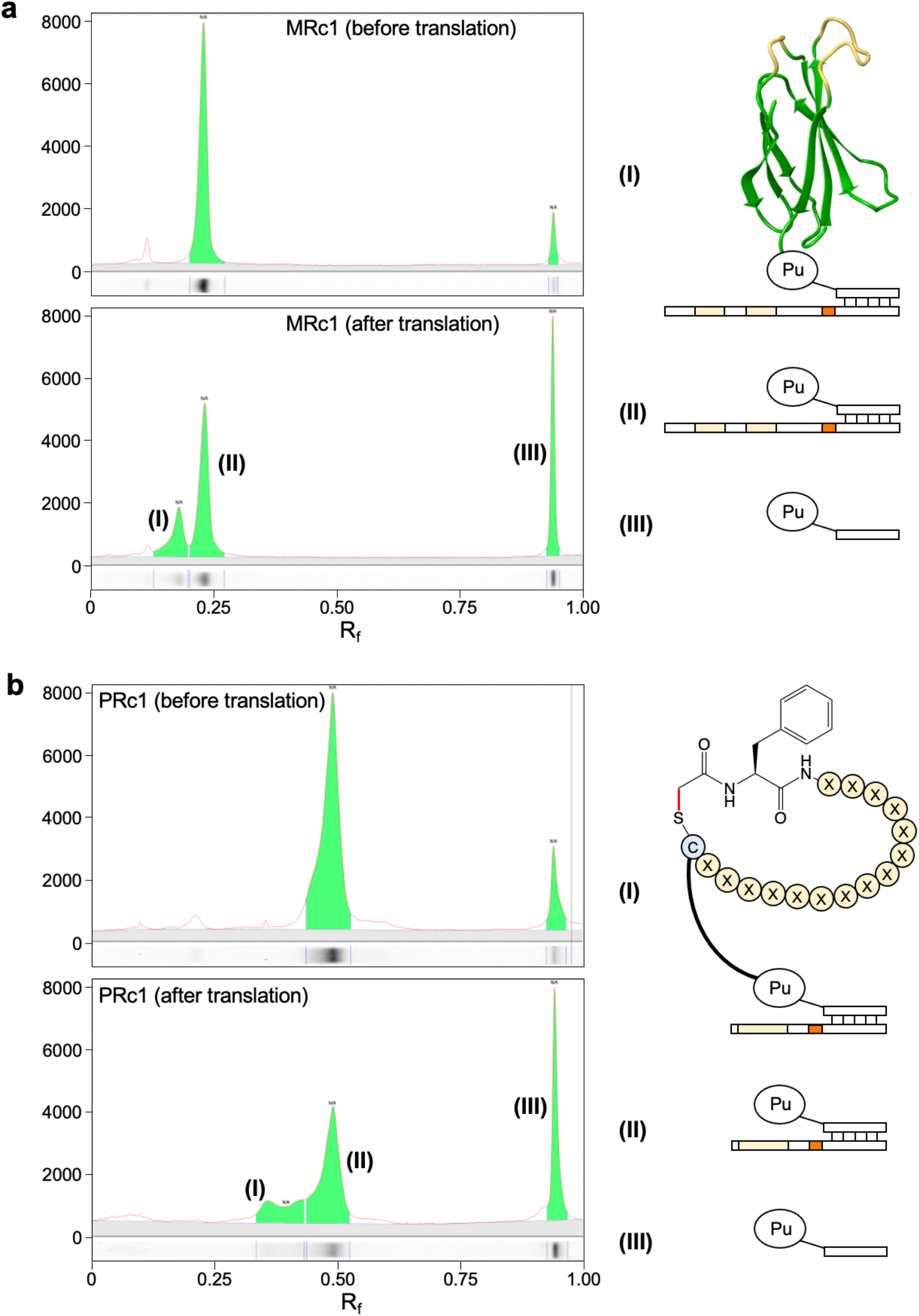
Raw data for determining puromycin conjugation efficiencies of mRNA libraries carrying monobodies or macrocyclic peptides. The efficiencies were calculated by I/(I + II + III).

**Table S1.** Oligonucleotides for preparation of NGS samples. * Duplicate experiment for Fig. 1c.

| Template |  |  | Forward primer | Reverse primer |
| --- | --- | --- | --- | --- |
| AUG-NNW<sub>3</sub><br>libraries | SD8-fibronectin | *pull-down*<sup>(-)</sup> | 50 μM SD8barcode1 | 12.5 μM HATagR23+3<br>12.5 μM HATagR23+2<br>12.5 μM HATagR23+1<br>12.5 μM HATagR23 mix |
|  |  | *pull-down*<sup>(+)</sup> | 50 μM SD8barcode2 |  |
|  | *SD8-fibronectin | *pull-down*<sup>(-)</sup> | 50 μM SD8barcode3 |  |
|  |  | *pull-down*<sup>(+)</sup> | 50 μM SD8barcode4 |  |
|  | SD8-lipocalin | *pull-down*<sup>(-)</sup> | 50 μM SD8barcode5 |  |
|  |  | *pull-down*<sup>(+)</sup> | 50 μM SD8barcode6 |  |
|  | SDcat-fibronectin | *pull-down*<sup>(-)</sup> | 50 μM SDcatbarcode9 |  |
|  |  | *pull-down*<sup>(+)</sup> | 50 μM SDcatbarcode10 |  |
| VVN<sub>2</sub>-<br>UAG-Z<br>libraries | Fibronectin-UAG | *pull-down*<sup>(-)</sup> | 25 μM MonoMidF22 | 50 μM an21-3barcode11 |
|  |  | *pull-down*<sup>(+)</sup> | 25 μM MonoMidF22+2 mix | 50 μM an21-3barcode12 |
|  | Lipocalin-UAG | *pull-down*<sup>(-)</sup> | 25 μM LcnMidF22 | 50 μM an21-3barcode13 |
|  |  | *pull-down*<sup>(+)</sup> | 25 μM LcnMidF22+2 mix | 50 μM an21-3barcode14 |

**Table S2.**
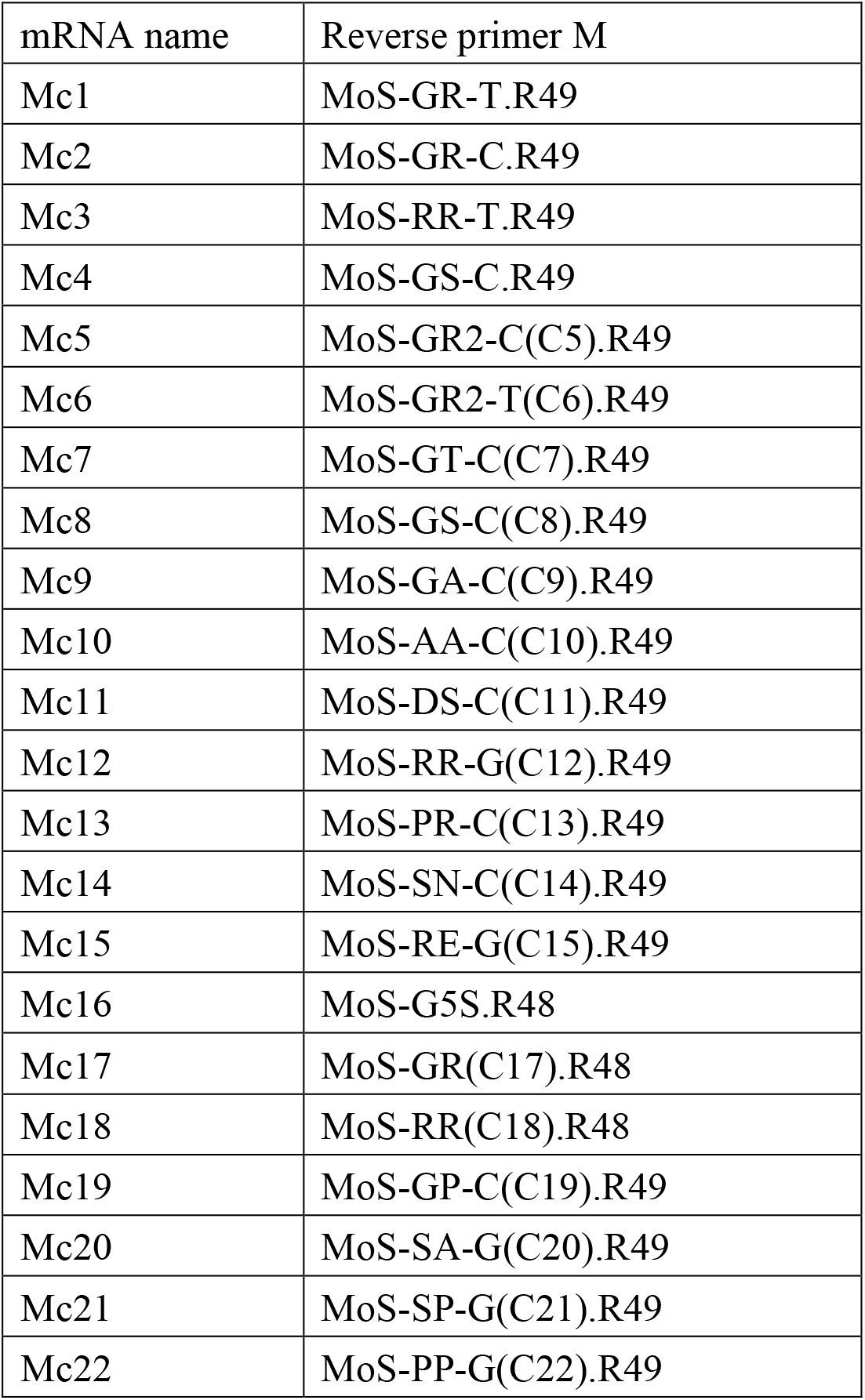
Reverse primers for the preparation of mRNAs (Mc1-Mc22).

| mRNA name | Reverse primer M |
| --- | --- |
| Mc1 | MoS-GR-T.R49 |
| Mc2 | MoS-GR-C.R49 |
| Mc3 | MoS-RR-T.R49 |
| Mc4 | MoS-GS-C.R49 |
| Mc5 | MoS-GR2-C(C5).R49 |
| Mc6 | MoS-GR2-T(C6).R49 |
| Mc7 | MoS-GT-C(C7).R49 |
| Mc8 | MoS-GS-C(C8).R49 |
| Mc9 | MoS-GA-C(C9).R49 |
| Mc10 | MoS-AA-C(C10).R49 |
| Mc11 | MoS-DS-C(C11).R49 |
| Mc12 | MoS-RR-G(C12).R49 |
| Mc13 | MoS-PR-C(C13).R49 |
| Mc14 | MoS-SN-C(C14).R49 |
| Mc15 | MoS-RE-G(C15).R49 |
| Mc16 | MoS-G5S.R48 |
| Mc17 | MoS-GR(C17).R48 |
| Mc18 | MoS-RR(C18).R48 |
| Mc19 | MoS-GP-C(C19).R49 |
| Mc20 | MoS-SA-G(C20).R49 |
| Mc21 | MoS-SP-G(C21).R49 |
| Mc22 | MoS-PP-G(C22).R49 |

**Table S3.** Reverse primers for the preparation of mRNA libraries (MRc1, MRc13, and MRc16).

| mRNA name | Reverse primer MR |
| --- | --- |
| MRc1 | MoS-GR-T.R49 |
| MRc13 | MoS-PR-C(C13).R49 |
| MRc16 | MoS-G5S.R48 |

**Table S4.** Reverse primers for the preparation of mRNA libraries (PRc1, PRc13, and PRc16).

| mRNA name | Reverse primer PR |
| --- | --- |
| PRc1 | Peptide-GR-T.R55 |
| PRc13 | Peptide-PR-C.R55 |
| PRc16 | Peptide-G5S.R54 |

## References

1 Voges, D., Watzele, M., Nemetz, C., Wizemann, S. & Buchberger, B. Analyzing and enhancing mRNA translational efficiency in an Escherichia coli in vitro expression system. Biochem. Biophys. Res. Commun. 318, 601–614 (2004).

2 Allert, M., Cox, J. C. & Hellinga, H. W. Multifactorial determinants of protein expression in prokaryotic open reading frames. J. Mol. Biol. 402, 905–918 (2010).

3 Tuller, T., Waldman, Y. Y., Kupiec, M. & Ruppin, E. Translation efficiency is determined by both codon bias and folding energy. Proc. Natl. Acad. Sci. U.S.A. 107, 3645–3650 (2010).

4 Espah Borujeni, A. et al. Precise quantification of translation inhibition by mRNA structures that overlap with the ribosomal footprint in N-terminal coding sequences. Nucleic Acids Res. 45, 5437–5448 (2017).

5 Qing, G., Xia, B. & Inouye, M. Enhancement of translation initiation by A/T-Rich sequences downstream of the initiation codon in Escherichia coli. J. Mol. Microbiol. Biotechnol. 6, 133–144 (2003).

6 Mitarai, N., Sneppen, K. & Pedersen, S. Ribosome collisions and translation efficiency: optimization by codon usage and mRNA destabilization. J. Mol. Biol. 382, 236–245 (2008).

7 Tuller, T. et al. An evolutionarily conserved mechanism for controlling the efficiency of protein translation. Cell 141, 344–354 (2010).

8 Tuller, T. et al. Composite effects of gene determinants on the translation speed and density of ribosomes. Genome Biol. 12, R110 (2011).

9 Kudla, G., Murray, A. W., Tollervey, D. & Plotkin, J. B. Coding-sequence determinants of gene expression in Escherichia coli. Science 324, 255–258 (2009).

10 Goodman, D. B., Church, G. M. & Kosuri, S. Causes and effects of N-terminal codon bias in bacterial genes. Science 342, 475–479 (2013).

11 Cambray, G., Guimaraes, J. C. & Arkin, A. P. Evaluation of 244,000 synthetic sequences reveals design principles to optimize translation in Escherichia coli. Nat. Biotechnol. 36, 1005–1015 (2018).

12 Verma, M. et al. A short translational ramp determines the efficiency of protein synthesis. Nat. Commun. 10, 5774 (2019).

13 Osterman, I. A. et al. Translation at first sight: the influence of leading codons. Nucleic Acids Res. 48, 6931–6942 (2020).

14 Geigenmüller, U. & Nierhaus, K. H. Significance of the third tRNA binding site, the E site, on E. coli ribosomes for the accuracy of translation: an occupied E site prevents the binding of non-cognate aminoacyl-tRNA to the A site. EMBO J. 9, 4527–4533 (1990).

15 Nierhaus, K. H. Decoding errors and the involvement of the E-site. Biochimie 88, 1013–1019 (2006).

16 Zaher, H. S. & Green, R. Fidelity at the molecular level: lessons from protein synthesis. Cell 136, 746–762 (2009).

17 Schilling-Bartetzko, S., Bartetzko, A. & Nierhaus, K. H. Kinetic and thermodynamic parameters for tRNA binding to the ribosome and for the translocation reaction. J. Biol. Chem. 267, 4703–4712 (1992).

18 Mottagui-Tabar, S., Björnsson, A. & Isaksson, L. A. The second to last amino acid in the nascent peptide as a codon context determinant. EMBO J. 13, 249–257 (1994).

19 Shimizu, Y. et al. Cell-free translation reconstituted with purified components. Nat. Biotechnol. 19, 751–755 (2001).

20 Ohashi, H., Shimizu, Y., Ying, B. W. & Ueda, T. Efficient protein selection based on ribosome display system with purified components. Biochem. Biophys. Res. Commun. 352, 270–276 (2007).

21 Reid, P. C., Goto, Y., Katoh, T. & Suga, H. Charging of tRNAs using ribozymes and selection of cyclic peptides containing thioethers. Methods Mol. Biol. 805, 335–348 (2012).

22 Kawakami, T., Ishizawa, T. & Murakami, H. Extensive reprogramming of the genetic code for genetically encoded synthesis of highly N-alkylated polycyclic peptidomimetics. J. Am. Chem. Soc. 135, 12297–12304 (2013).

23 Kondo, T. et al. Antibody-like proteins that capture and neutralize SARS-CoV-2. Sci. Adv. 6, 12 (2020).

24 Roberts, R. W. & Szostak, J. W. RNA-peptide fusions for the in vitro selection of peptides and proteins. Proc. Natl. Acad. Sci. U.S.A. 94, 12297–12302 (1997).

25 Nemoto, N., Miyamoto-Sato, E., Husimi, Y. & Yanagawa, H. In vitro virus: Bonding of mRNA bearing puromycin at the 3’-terminal end to the C-terminal end of its encoded protein on the ribosome in vitro. FEBS Lett. 414, 405–408 (1997).

26 Ishizawa, T., Kawakami, T., Reid, P. C. & Murakami, H. TRAP Display: A high-speed selection method for the generation of functional polypeptides. J. Am. Chem. Soc. 135, 5433–5440 (2013).

27 Koide, A., Bailey, C. W., Huang, X. & Koide, S. The fibronectin type III domain as a scaffold for novel binding proteins. J. Mol. Biol. 284, 1141–1151 (1998).

28 Beste, G., Schmidt, F. S., Stibora, T. & Skerra, A. Small antibody-like proteins with prescribed ligand specificities derived from the lipocalin fold. Proc. Natl. Acad. Sci. U.S.A. 96, 1898–1903 (1999).

29 Gebauer, M., Schiefner, A., Matschiner, G. & Skerra, A. Combinatorial design of an Anticalin directed against the extra-domain b for the specific targeting of oncofetal fibronectin. J. Mol. Biol. 425, 780–802 (2013).

30 Ude, S. et al. Translation elongation factor EF-P alleviates ribosome stalling at polyproline stretches. Science 339, 82–85 (2013).

31 Doerfel, L. K. et al. EF-P is essential for rapid synthesis of proteins containing consecutive proline residues. Science 339, 85–88 (2013).

32 Doerfel, L. K. et al. Entropic contribution of elongation factor P to proline positioning at the catalytic center of the ribosome. J. Am. Chem. Soc. 137, 12997–13006 (2015).

33 Koutmou, K. S. et al. Ribosomes slide on lysine-encoding homopolymeric A stretches. Elife 4 (2015).

34 Lorenz, R. et al. ViennaRNA Package 2.0. Algorithms Mol. Biol. 6, 26 (2011).

35 Zaher, H. S. & Green, R. Quality control by the ribosome following peptide bond formation. Nature 457, 161–166 (2009).

36 Murakami, H., Saito, H. & Suga, H. A versatile tRNA aminoacylation catalyst based on RNA. Chem. Biol. 10, 655–662 (2003).

37 Murakami, H., Ohta, A., Ashigai, H. & Suga, H. A highly flexible tRNA acylation method for non-natural polypeptide synthesis. Nat. Methods 3, 357–359 (2006).

38 Ohuchi, M., Murakami, H. & Suga, H. The flexizyme system: a highly flexible tRNA aminoacylation tool for the translation apparatus. Curr. Opin. Chem. Biol. 11, 537–542 (2007).

39 Kondo, T. et al. cDNA TRAP display for rapid and stable in vitro selection of antibody-like proteins. Chem. Commun. 57, 2416–2419 (2021).

40 Bossi, L. & Roth, J. R. The influence of codon context on genetic-code translation. Nature 286, 123–127 (1980).

41 Bossi, L. Context effects: translation of UAG codon by suppressor tRNA is affected by the sequence following UAG in the message. J. Mol. Biol. 164, 73–87 (1983).

42 Miller, J. H. & Albertini, A. M. Effects of surrounding sequence on the suppression of nonsense codons. J. Mol. Biol. 164, 59–71 (1983).

43 Smolskaya, S., Zhang, Z. J. & Alfonta, L. Enhanced yield of recombinant proteins with site-specifically incorporated unnatural amino acids using a cell-free expression system. PLoS One 8 (2013).

44 Xu, H. et al. Re-exploration of the codon context effect on amber codon-guided incorporation of noncanonical amino acids in Escherichia coli by the blue-white screening assay. Chembiochem 17, 1250–1256 (2016).

45 Grosjean, H., Söll, D. G. & Crothers, D. M. Studies of the complex between transfer RNAs with complementary anticodons. I. Origins of enhanced affinity between complementary triplets. J. Mol. Biol. 103, 499–519 (1976).

46 Engelberg-Kulka, H. UGA suppression by normal tRNA Trp in Escherichia coli: codon context effects. Nucleic Acids Res. 9, 983–991 (1981).

47 Nilsson, M. & Rydén-Aulin, M. Glutamine is incorporated at the nonsense codons UAG and UAA in a suppressor-free Escherichia coli strain. Biochim. Biophys. Acta 1627, 1–6 (2003).

48 Murphy, F. V. T. & Ramakrishnan, V. Structure of a purine-purine wobble base pair in the decoding center of the ribosome. Nat. Struct. Mol. Biol. 11, 1251–1252 (2004).

49 Letzring, D. P., Dean, K. M. & Grayhack, E. J. Control of translation efficiency in yeast by codon-anticodon interactions. RNA 16, 2516–2528 (2010).

50 Gamble, C. E., Brule, C. E., Dean, K. M., Fields, S. & Grayhack, E. J. Adjacent codons act in concert to modulate translation efficiency in yeast. Cell 166, 679–690 (2016).

51 Tunney, R. et al. Accurate design of translational output by a neural network model of ribosome distribution. Nat. Struct. Mol. Biol. 25, 577–582 (2018).

52 Menninger, J. R. Peptidyl transfer RNA dissociates during protein synthesis from ribosomes of Escherichia coli. J. Biol. Chem. 251, 3392–3398 (1976).

53 Fahlman, R. P., Dale, T. & Uhlenbeck, O. C. Uniform binding of aminoacylated transfer RNAs to the ribosomal A and P sites. Mol. Cell 16, 799–805 (2004).

54 Olejniczak, M., Dale, T., Fahlman, R. P. & Uhlenbeck, O. C. Idiosyncratic tuning of tRNAs to achieve uniform ribosome binding. Nat. Struct. Mol. Biol. 12, 788–793 (2005).

55 Yamagishi, Y. et al. Natural product-like macrocyclic N-methyl-peptide inhibitors against a ubiquitin ligase uncovered from a ribosome-expressed de novo library. Chem. Biol. 18, 1562–1570 (2011).

56 Kawakami, T. et al. In vitro selection of multiple libraries created by genetic code reprogramming to discover macrocyclic peptides that antagonize VEGFR2 activity in living cells. ACS Chem. Biol. 8, 1205–1214 (2013).

57 Huang, Y., Wiedmann, M. M. & Suga, H. RNA display methods for the discovery of bioactive macrocycles. Chem. Rev. 119, 10360–10391 (2019).

58 Kawakami, T. & Murakami, H. Genetically encoded libraries of nonstandard peptides. J. Nucleic Acids 2012, 713510 (2012).

59 Goto, Y. et al. Reprogramming the translation initiation for the synthesis of physiologically stable cyclic peptides. ACS Chem. Biol. 3, 120–129 (2008).

60 Nagumo, Y., Fujiwara, K., Horisawa, K., Yanagawa, H. & Doi, N. PURE mRNA display for in vitro selection of single-chain antibodies. J. Biochem. 159, 519–526 (2016).

61 Takahashi, K., Sunohara, M., Terai, T., Kumachi, S. & Nemoto, N. Enhanced mRNA-protein fusion efficiency of a single-domain antibody by selection of mRNA display with additional random sequences in the terminal translated regions. Biophys. Physicobiol. 14, 23–28 (2017).

62 Tajima, K., Katoh, T. & Suga, H. Drop-off-reinitiation triggered by EF-G-driven mistranslocation and its alleviation by EF-P. Nucleic Acids Res. 50, 2736–2753 (2022).

63 Kelsic, E. D. et al. RNA structural determinants of optimal codons revealed by MAGE-Seq. Cell Syst. 3, 563-571.e566 (2016).

